# A parasitic fungus employs mutated eIF4A to survive on rocaglate-synthesizing *Aglaia* plants

**DOI:** 10.1101/2022.07.04.498659

**Authors:** Mingming Chen, Naoyoshi Kumakura, Ryan Muller, Yuichi Shichino, Madoka Nishimoto, Mari Mito, Pamela Gan, Nicholas T. Ingolia, Ken Shirasu, Takuhiro Ito, Shintaro Iwasaki

## Abstract

Plants often generate secondary metabolites as defense mechanisms against parasites. Although some fungi may potentially overcome the barrier of antimicrobial compounds, only a limited number of examples and molecular mechanisms of resistance have been reported. Here, we found an *Aglaia* plant-parasitizing fungus that overcomes the toxicity of rocalgates, which are translation inhibitors synthesized by the plant, through an amino acid substitution in a translation initiation factor (eIF). *De novo* transcriptome assembly revealed that the fungus belongs to *Ophiocordyceps* genus and its eIF4A, a molecular target of rocaglates, contains a amino acid substitution critical for rocaglate binding. Ribosome profiling harnessing a cucumber-infecting fungus, *Colletotrichum orbiculare*, demonstrated that the translational inhibitory effects of rocaglates were largely attenuated by the mutation found in the *Aglaia* parasite. The engineered *Colletotrichum orbiculare* showed a survival advantage on cucumber plants with rocaglates. Our study exemplifies a plant-fungus tug-of-war centered on secondary metabolites produced by host plants.

## Introduction

Fungi that infect plants are of great economic relevance because they cause severe crop losses (∼10%) worldwide (Oerke, 2006). Therefore, the mechanisms underlying plant-fungus interactions have attracted great interest and have been extensively studied (Lo Presti et al., 2015). Secondary metabolites with antimicrobial activities are among the means naturally developed by plants for the control of fungal infections (Collemare et al., 2019). For example, tomatine, a glycoalkaloid secreted from the leaves and stems of tomato, has both fungicidal properties and insecticidal activities (Vance et al., 1987). Camalexin, an indole alkaloid produced by Brassicaceae plants, including the model plant *Arabidopsis thaliana,* also has antifungal properties (Nafisi et al., 2007).

However, some fungi can overcome these toxic compounds to infect plants. The best-known strategy is the detoxification of antifungal compounds by the secretion of specific enzymes (Crombie et al., 1986; Osbourn et al., 1995; Pareja-Jaime et al., 2008). Thus, plants and infectious fungi are engaged in an arms race during the course of evolution. However, other than detoxification, the mechanistic diversity of the plant-fungus competition centered on plant secondary metabolites is largely unknown.

Rocaglates, small molecules synthesized in plants of the genus *Aglaia*, exemplify antifungal secondary metabolites (Engelmeier et al., 2000; Iyer et al., 2020). In addition to its antifungal properties, this group of compounds is of particular interest because of its antitumor activities (Alachkar et al., 2013; Bordeleau et al., 2008; Cencic et al., 2009; Chan et al., 2019; Ernst et al., 2020; Lucas et al., 2009; Manier et al., 2017; Nishida et al., 2021; Santagata et al., 2013; Skofler et al., 2021; Thompson et al., 2021, 2017; Wilmore et al., 2021; Wolfe et al., 2014). Moreover, recent studies have suggested potency against viruses including SARS-CoV-2 (Müller et al., 2021, 2020). Rocaglates target translation initiation factor (eIF) 4A, a DEAD-box RNA binding protein, and function as potent translation inhibitors with a unique mechanism: rocaglate treatment does not phenocopy the loss-of-function of eIF4A but instead leads to gain-of-function. Although eIF4A activates the translation of cellular mRNA through ATP-dependent RNA binding, rocaglates impose polypurine (A and G repeated) sequence selectivity on eIF4A, bypassing the ATP requirements and evoking mRNA-selective translation repression (Chen et al., 2021; Chu et al., 2020, 2019; Iwasaki et al., 2019, 2016; Rubio et al., 2014; Wolfe et al., 2014). The artificial anchoring of eIF4A 1) becomes a steric hindrance to scanning 40S ribosomes (Iwasaki et al., 2019, 2016), 2) masks cap structure of mRNA by tethering eIF4F (Chu et al., 2020), and 3) reduces the available pool of eIF4A for active translation initiation events by the sequestration of eIF4A on mRNAs (Chu et al., 2020).

Since eIF4A is an essential gene for all eukaryotes, *Aglaia* plants must have a mechanism to evade the cytotoxicity of the rocaglates they produce. This self-resistance is achieved by the unique amino acid substitutions at the sites in eIF4A proteins where rocaglates directly associate (Iwasaki et al., 2019). Given the high evolutionary conservation of eIF4A and thus the rocaglate binding pocket (Iwasaki et al., 2019), the compounds may target a wide array of natural fungi.

Irrespective of the antifungal nature of rocaglates, we found a parasitic fungus able to grow on *Aglaia* plants. *De novo* transcriptome analysis from the fungus revealed that this species belongs to the *Ophiocordyceps* genus, which is well known to infect ants and cause a “zombie” phenotype (Andersen et al., 2009; Araújo and Hughes, 2019; de Bekker et al., 2017), but forms a distinct branch in the taxon. Strikingly, eIF4A from this fungus possessed an amino acid substitution in the rocaglate binding site and thus showed resistance to the compound. Using *Colletotrichum orbiculare*, a cucumber-infecting fungus, as a model, we demonstrated that the genetically engineered fungus with the substitution showed insensitivity to the translation repression evoked by rocaglates, facilitating its infection of plants even in the presence of this compound. Our results indicate fungal resistance to plant secondary metabolites independent of detoxification enzymes and a unique contest between plants and fungi centered on secondary metabolites synthesized in the host plant.

## Results

### Identification of a fungal parasite on the rocaglate-producing plant Aglaia

Considering that *Aglaia* plants possess antifungal rocaglates (Engelmeier et al., 2000; Iyer et al., 2020), parasitic fungi should have difficulty infecting rocaglate-producing plants. In contrast to this idea, we identified a fungus growing on the surface of the stem of the *Aglaia odorata* plant with tremendous vitality (Figure 1A). To characterize this fungus, we isolated the RNA, conducted RNA sequencing (RNA-Seq), reconstructed the transcriptome, and annotated the functionality of each gene (Supplementary Table 1).

**Figure 1.**
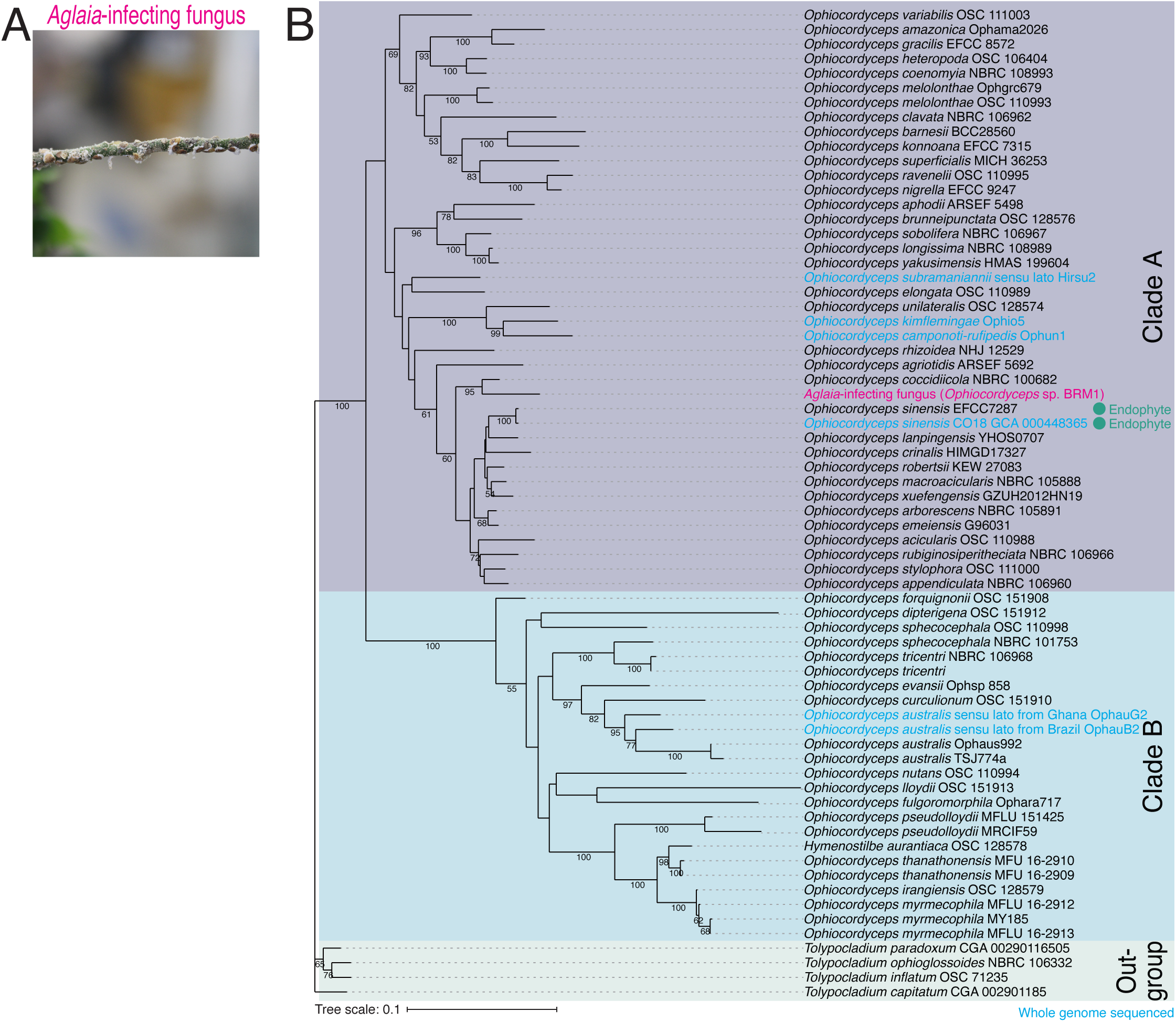
Identification of *Aglaia*-parasitic *Ophiocordyceps* sp. BRM1. (A) Image of a parasite fungus growing on *Aglaia odorata*. (B) Multilocus phylogenetic tree of *Ophiocordyceps* species generated from maximum-likelihood phylogenetic analysis of ITS, SSU, LSU, *RPB1*, and *TEF1a* sequences. *Tolypocladium* species were used as outgroups. The best DNA substitution models of ITS, LSU, SSU, *RPB1* and *TEF1α* were calculated as TIM3ef+G4, TIM1+I+G4, TIM3ef+I+G4, TrN+I+G4, and TIM1+I+G4, respectively. Numbers on branches are percent support values out of 1,000 bootstrap replicates. Only bootstrap values greater than 50% support are shown. Endophytes are highlighted with green dots.

This *Aglaia*-infecting fungus belonged to the *Ophiocordyceps* genus, which is known as zombie-ant fungus (Andersen et al., 2009; Araújo and Hughes, 2019; de Bekker et al., 2017). *Ophiocordyceps* spp. belongs to the division Ascomycota and is one of the taxonomic group with the highest number of entomopathogenic species in all fungal genus. In the majority of cases, each of *Ophiocordyceps* spp. has its specific host insect species, develops fruiting bodies from remains of host insects, and produces spores. Although *Ophiocordyceps* spp. are known as insect pathogens, recent studies have detected a moth parasite *Ophiocordyceps sinensis* in many plant species, suggesting that *Ophiocordyceps* also has an endophytic lifestyle (Wang et al., 2020; Zhong et al., 2014). We performed BLASTn (Camacho et al., 2009) using the internal transcribed spacer (ITS) between rRNAs as a query and found that, among all the deposited nucleotide sequences in the database, 29 of the top 30 hits were from *Ophiocordyceps* species (Supplementary Table 2).

To identify the species-level taxon of the *Aglaia*-infecting fungus, we conducted a multilocus phylogenetic analysis for comparison with currently accepted species in the *Ophiocordyceps* genus (Supplementary Table 3). For this purpose, sequences of the ITS, small subunit ribosomal RNA (SSU), large subunit rRNA (LSU), translation elongation factor 1-alpha (*TEF1a*), and RNA polymerase II largest subunit (*RPB1*) were used as previously reported for the classification of *Ophiocordyceps* species(Xiao et al., 2017) (Supplementary Table 3). These sequences from 68 isolates were aligned, trimmed, and concatenated, resulting in a multiple sequence alignment comprising 3,910 nucleotide positions, including gaps (gene boundaries ITS, 1-463; LSU, 464-1,363; SSU, 1,364-2,248; *RPB1*, 2,249-2,922; *TEF1α*, 2,923-3,910). Then, the best-scoring maximum-likelihood (ML) tree was calculated from the concatenated sequence alignment using the selected DNA substitution models for each sequence (Figure 1B). This analysis indicated that the *Aglaia*-infecting fungus was distinct from the other species of *Ophiocordyceps* (Figure 1B). In particular, the strain isolated from *Aglaia* was positioned on a long branch separated from the most closely related strain, *Ophiocordyceps coccidiicola* NBRC 100682, was supported by the 95% bootstrap value. The separation of *Aglaia*-infecting fungus from other *Ophiocordyceps* species was also supported by single-locus alignments (Figure 1 — figure supplement 1), although the positions of the *Aglaia*-infecting fungus in the tree were different. We note that *RPB1*-locus alignment was an exception since the *de novo*-assembled transcriptome from the *Aglaia*-infecting fungus lacked the sequence of the homolog.

Given that the most closely related *Ophiocordyceps* species is sufficiently distinct from the *Aglaia*-infecting fungus in sequence and that no similar fungi grown in *Aglaia* plants were reported before, we named the fungus *Ophiocordyceps* sp. BRM1 (Berkeley, Ryan Muller, strain 1). Consistent with the isolation from *Aglaia* plant, this fungi was a close kin of *Ophiocordyceps* spp. known as endophytes (Figure 1B and Supplementary Table 3) (Wang et al., 2020; Zhong et al., 2014).

### Transcriptome assembly uncovers the unique mutation in eIF4A of the Aglaia-infecting fungus

The parasitic nature of *Ophiocordyceps* sp. BRM1 on plants producing the antifungal rocaglate led us to hypothesize that the fungus may have a mechanism to evade the toxicity of the compounds. Indeed, the host plant *Aglaia* achieves this task by introducing an amino acid substitution in eIF4A, a target of rocaglates (Iwasaki et al., 2019). The substituted amino acid (Phe163, amino acid position in human eIF4A1) lies at the critical interface for rocaglate interaction (Figure 2A) (Iwasaki et al., 2019). Accordingly, we investigated possible amino acid conversion in eIF4As of the *Ophiocordyceps* sp. BRM1. Among the *de novo*-assembled transcriptome, ∼60 DEAD-box RNA binding protein genes, including 4 isoforms of eIF4A homologs, were found (Supplementary Table 1).

**Figure 2.**
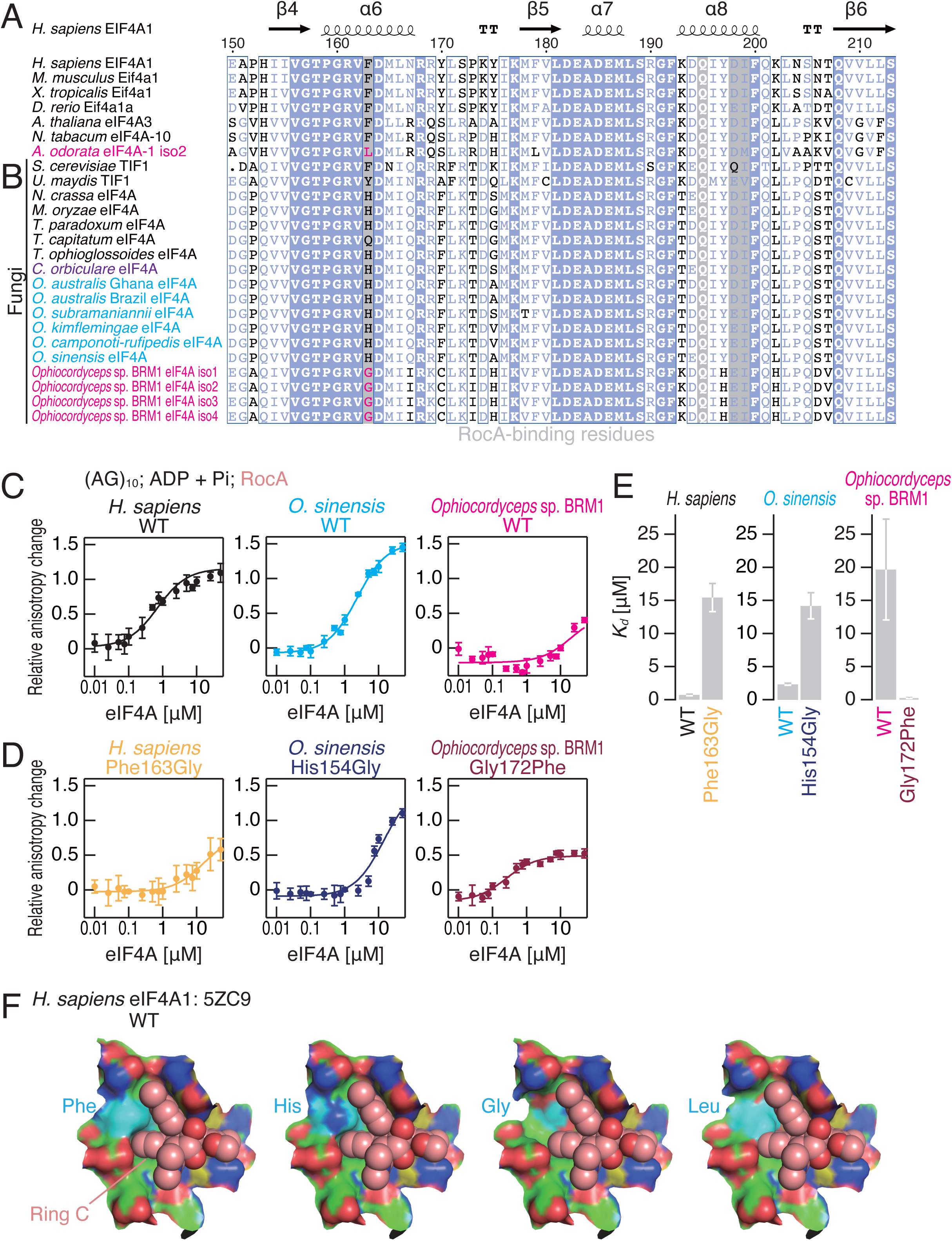
The effect of an amino acid substitution found in *Ophiocordyceps* sp. BRM1 eIF4A on RocA-mediated polypurine RNA clamping. (A and B) Alignments of eIF4A protein sequences from higher eukaryotes (A) and fungal species (B), including the *de novo*-assembled *Ophiocordyceps* sp. BRM1 eIF4A gene with four transcript isoforms (iso). (C and D) Fluorescence polarization assay for FAM-labeled RNA ([AG]_10_) (10 nM). WT (C) and mutated (D) eIF4A proteins from the indicated species were used. To measure ATP-independent RNA clamping induced by RocA (50 µM), ADP and Pi (1 mM each) were included in the reaction. Data represent the mean and s.d. (n = 3). (E) The summary of *K_d_* in C and D is depicted. Data represent the mean and s.d. (F) RocA (sphere model with light pink-colored carbon), modeled His, Gly, and Leu residues (surface model with cyan-colored carbon) at Phe163 residue in human eIF4A1 (surface model with green-colored carbon), and RNA (surface model with yellow-colored carbon) in the complex of human eIF4A1•RocA•AMPPNP•polypurine RNA (PDB: 5ZC9) (Iwasaki et al., 2019).

Strikingly, we observed amino acid conversion in the *Ophiocordyceps* sp. BRM1 eIF4A at the same residue as in the *Aglaia* plant eIF4A. Gly residues were found to replace Phe163 (human position) in all four transcript isoforms (from the same eIF4A gene) in *Ophiocordyceps* sp. BRM1 (Figure 2B and Figure 2 — figure supplement 1A), whereas His residues prevailed in the close kin of *Ophiocordyceps* species and other fungi.

### Gly153 in Ophiocordyceps sp. BRM1 eIF4A eliminated rocaglate-mediated polypurine RNA clamping

Indeed, we found that the Gly substitution confers rocaglate resistance on eIF4A. To investigate rocaglate-targetability, we harnessed the fluorescence polarization assay with fluorescein (FAM)-labeled short RNA and purified recombinant eIF4A proteins (Figure 2 — figure supplement 1B). As observed previously (Chen et al., 2021; Chu et al., 2020, 2019; Iwasaki et al., 2019, 2016; Naineni et al., 2021), rocaglamide A (RocA), a natural rocaglate derivative isolated from *Aglaia* plants (Figure 2 — figure supplement 1C) (Janprasert et al., 1992), clamped human eIF4A1 on polypurine RNA ([AG]_10_) in an ATP-independent manner (*e.g.*, in the presence of ADP + Pi) (Figure 2C left, 2E left and Table 1). Whereas substantially similar affinity to polypurine RNA was observed in eIF4A from *O. sinensis* (CO18 GCA 000448365) (Figure 2C middle, 2E middle and Table 1), the closest kin among whole-genome-sequenced *Ophiocordyceps* species (Figure 1B) (de Bekker et al., 2017), *Ophiocordyceps* sp. BRM1 eIF4A showed a fairly high *K_d_* (Figure 2C right, 2E right and Table 1).

**Table 1.**
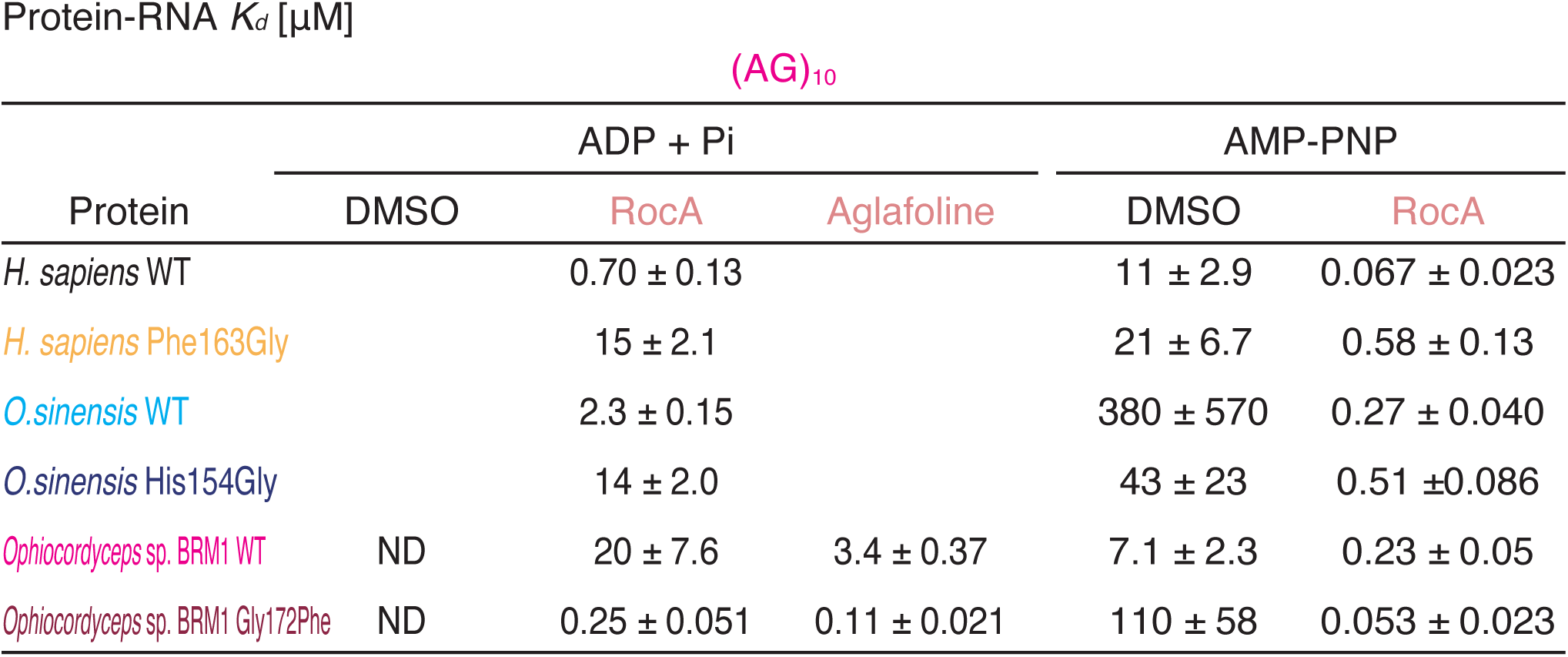
Summary of *K_d_* (µM) between eIF4A protein and RNAs. Fluorescence polarization assay between FAM-labeled RNA ([AG]_10_) and the indicated recombinant proteins was conducted to measure the *K_d_* in the presence of DMSO, RocA, or aglafoline.

Given the Phe163Gly substitution (human position) in *Ophiocordyceps* sp. BRM1 eIF4A (Figure 2A), we hypothesized that the amino acid substitution explains the differential sensitivity to RocA. Indeed, Gly to Phe substitution in the *Ophiocordyceps* sp. BRM1 eIF4A (Gly172Phe) sensitized the protein to RocA (Figure 2D right), reducing the *K_d_* significantly (Figure 2E right and Table 1). Conversely, the introduction of the Gly residue into human and *O. sinensis* eIF4As reduced the affinity to polypurine RNA (Figure 2D middle and left, 2E middle and left and Table 1).

A similar rocaglate sensitivity in RNA binding was also observed in AMP-PNP, an ATP ground state analog (Figure 2 — figure supplement 1D, 1E and Table 1). Unlike ADP + Pi, AMP-PNP allowed basal binding to polypurine RNAs in the absence of RocA. The affinity was further increased by RocA in eIF4A with Phe or His residues at 163 (human position). In contrast, the Gly residue strongly suppressed the affinity changes (Figure 2 — figure supplement 1D, 1E and Table 1).

Taking these biochemical data together, we concluded that the *Ophiocordyceps* sp. BRM1 eIF4A evades rocaglate targeting by substituting a critical amino acid involved in its binding. When Phe163 was replaced by Gly in the crystal structure of the human eIF4A1•RocA complex (Iwasaki et al., 2019), the π-π stacking with ring C of RocA was totally lost (Figure 2F and Figure 2 — figure supplement 1C), likely leading to reduced affinity for RocA. This mechanism to desensitize eIF4A to rocaglates was distinct from the Leu substitution found in *Aglaia*, which fills the space of the rocaglate binding pocket and thus prevents the interaction (Iwasaki et al., 2019).

Our data showed that eIF4A with His at 163 (human position) is also a target of rocaglate (Figure 2C-E and Figure 2 — figure supplement 1D, 1E). This is most likely due to the functional replacement of the aromatic ring in Phe by the imidazole ring in His for stacking with ring C of rocaglates (Figure 2F). This suggested that a wide array of fungi that possess the His variant (Figure 2A), including *C. orbiculare* (see below for details), are also susceptible to rocaglates.

### Gly153 found in Ophiocordyceps *sp. BRM1* eIF4A confers resistance to rocaglate-induced translational repression

The reduced affinity to polypurine RNA gained in the *Ophiocordyceps* sp. BRM1 by Gly substitution led us to investigate the impact on rocaglate-mediated translation repression. To test this, we applied a reconstituted translation system with human factors (Iwasaki et al., 2019; Machida et al., 2018; Yokoyama et al., 2019). As observed in an earlier study (Iwasaki et al., 2019), this system enabled the recapitulation of translation reduction from polypurine motif-possessing reporter mRNA in a RocA dose-dependent manner (Figure 3A and Figure 3 — figure supplement 1A). In contrast, replacing wild-type human eIF4A1 with the Phe163Gly mutant prevented translation reduction by RocA (Figure 3A), which was consistent with the affinity between the recombinant human eIF4A1 proteins and polypurine RNA (Figures 2 and Figure 2 — figure supplement 1).

**Figure 3.**
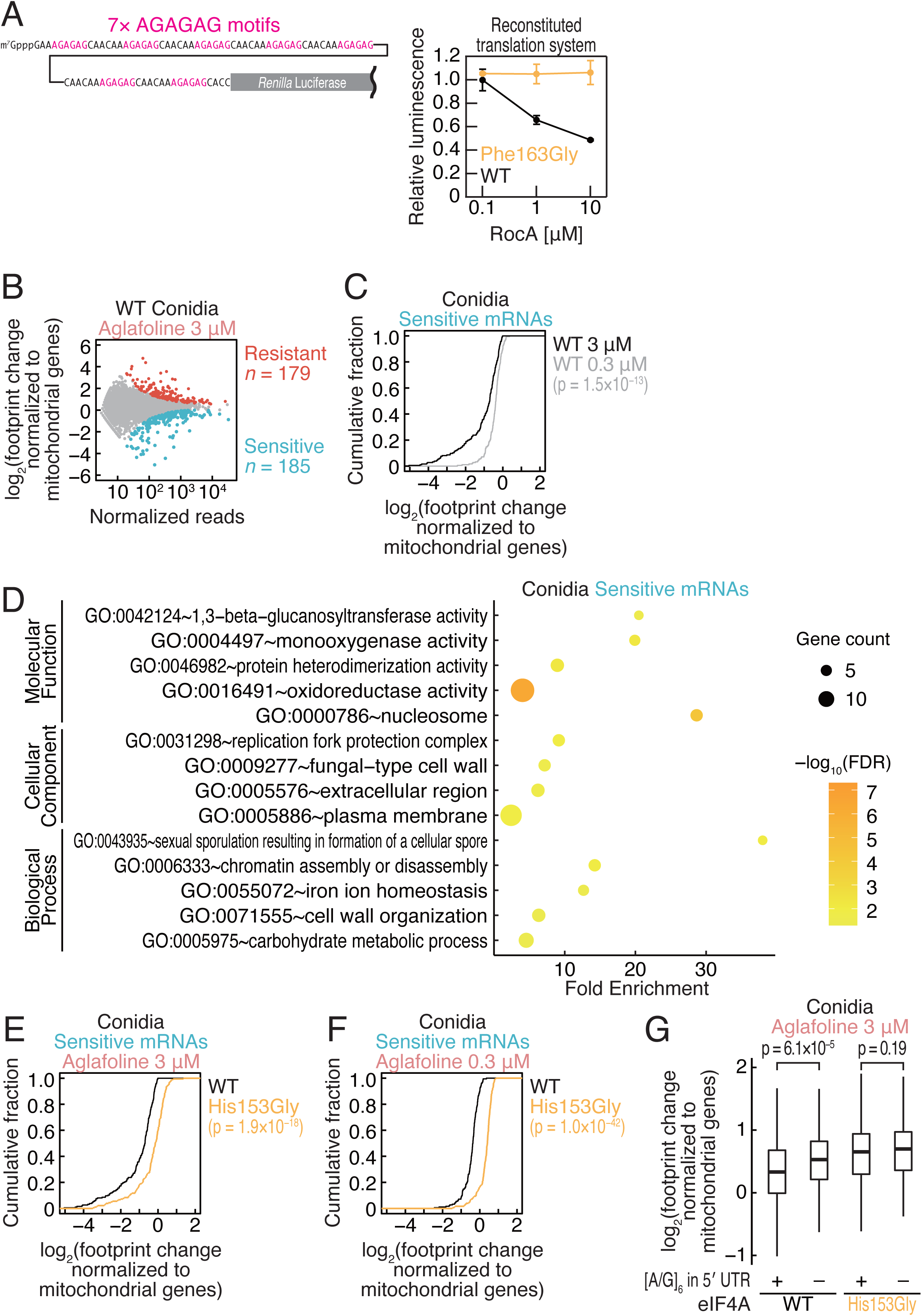
The amino acid substitution in the *Ophiocordyceps* sp. BRM1 eIF4A confers translational resistance to rocaglates in fungi. (A) RocA-mediated translational repression recapitulated by an *in vitro* reconstitution system with human factors. Recombinant proteins of *H. sapiens* eIF4A1 WT or Phe163Gly were added to the reaction with RocA. Reporter mRNA with polypurine motifs (left) was translated in the reaction. (B) MA (M, log ratio; A, mean average) plot of ribosome footprint changes caused by 3 µM aglafoline treatment in *C. orbiculare* eIF4A^WT^ conidia. Resistant and sensitive mRNAs (false discovery rate [FDR] < 0.05) are highlighted. Data were normalized to mitochondrial footprints, which were used as internal spike-ins (Iwasaki et al., 2016). (C) Cumulative distribution of the ribosome footprint changes for aglafoline-sensitive mRNAs (defined in B) in *C. orbiculare* eIF4A^WT^ conidia with 0.3 or 3 µM aglafoline treatment. (D) Gene Ontology (GO) term analysis for aglafoline-sensitive mRNAs (defined in B). GO terms associated with yeast homologs were analyzed by DAVID (Huang et al., 2009a, 2009b). (E and F) Cumulative distribution of the ribosome footprint changes for aglafoline-sensitive mRNAs (defined in B) by 3 µM (E) and 0.3 µM (F) aglafoline treatment in *C. orbiculare* eIF4A^WT^ and eIF4A^His153Gly^ conidia. (G) Box plot of ribosome footprint changes caused by 3 µM aglafoline treatment in conidia across mRNAs with or without [A/G]_6_ motif in 5′ UTR. The p values in C, E, F, and G were calculated by the Mann–Whitney *U* test.

We further tested the impact of the Gly conversion in eIF4A in a fungus. Due to the difficulty of culturing and manipulating the genetics of *Ophiocordyceps* sp. BRM1 (data not shown), we instead harnessed *Colletotrichum orbiculare*, an anthracnose-causing fungus (Gan et al., 2019, 2013). Through homology-directed repair induced by CRISPR–Cas9-mediated genome cleavage, we replaced endogenous eIF4A with wild-type (WT) or Gly-mutated (His153Gly) *C. orbiculare* eIF4A (Figure 2 — figure supplement 1A and Figure 3 — figure supplement 1B and 1C).

Since the culture of the isolated strains requires a significant amount of the compounds, we used aglafoline (methyl rocaglate) (Figure 2 — figure supplement 1C), a less expensive, commercially available natural derivative of rocaglates (Ko et al., 1992), instead of RocA. The difference between RocA and aglafoline was the dimethylamide group and methoxycarbonyl group (Figure 2 — figure supplement 1C), which do not contribute to association with eIF4A or polypurine RNA (Iwasaki et al., 2019), suggesting that the compounds should have similar mechanisms of action. As expected, aglafoline resulted in essentially the same molecular phenotype of ATP-independent polypurine clamping of the *Ophiocordyceps* sp. BRM1 eIF4A (Figure 3 — figure supplement 1D, 1E and Table 1) as RocA (Figure 2C-E and Table 1).

To understand the translation repression induced by aglafoline in a genome-wide manner, we applied ribosome profiling, a technique based on deep sequencing of ribosome-protected RNA fragments (*i.e.*, ribosome footprints) generated by RNase treatment (Ingolia et al., 2019, 2009; Iwasaki and Ingolia, 2017), to the isolated fungus strains. The ribosome footprints obtained from *C. orbiculare* (in the culture of conidia and mycelia) showed the signatures of this experiment: two peaks of footprint length at ∼22 nt and ∼30 nt (Figure 3 — figure supplement 2A), which respectively represent the absence or presence of A-site tRNA in the ribosome (Lareau et al., 2014; Wu et al., 2019), and 3-nt periodicity along the open reading frame (ORF) (Figure 3 — figure supplement 2B and 2C).

Strikingly, ribosome profiling revealed that His153Gly confers translational resistance to aglafoline in *C. orbiculare*. Consistent with the mRNA-selective action of the compound, we observed that a subset of mRNAs showed high aglafoline sensitivity in protein synthesis (Figure 3B), and the reduction in translation in conidia was compound-dose dependent (Figure 3C). Intriguingly, we observed that genes associated with carbohydrate metabolism and cell wall organization, which are both essential processes for fungal cell survival, were susceptible to translation repression by rocaglate (Figure 3D). The suppression of protein synthesis from aglafoline-sensitive mRNAs was attenuated by His153Gly substitution at both high (3 µM) and low (0.3 µM) concentrations of aglafoline (Figure 3E and 3F). Consistent with the earlier reports (Chen et al., 2021; Chu et al., 2020, 2019; Iwasaki et al., 2019, 2016), the overall reduction in protein synthesis associated with the presence of polypurine motifs in 5′ UTR (Figure 3G). However, His153Gly substitution compromised the polypurine dependent translation repression.

Although genetic engineering conferred similar aglafoline-insensitive translation in mycelia (Figure 3 — figure supplement 3A-E), the aglafoline-sensitive mRNAs were distinct from those in conidia (Figure 3 — figure supplement 3F). This was due to the exclusive expression of those sensitive mRNAs in each cell state (Figure 3 — figure supplement 3G), suggesting differential impacts of the rocaglates during the fungal life cycle.

### Rocaglate-resistant fungi show an advantage in infection of plants with rocaglates

We were intrigued to test the role of Gly substitution in the parasitic property of fungi. Here, we used the infection process of *C. orbiculare* on cucumber leaves as a model system. The conidia of WT or His153Gly eIF4A-recombined strains were sprayed on *Cucumis sativus* (cucumber) cotyledons, and the biomass after inoculation with rocaglate was quantified (Figure 4A). Indeed, aglafoline reduced the biomass of the WT eIF4A-recombined strain on cucumber leaves, showing the antifungal effect of rocaglate (Figure 4B). In stark contrast, the His153Gly mutation in eIF4A affected fungal growth on cucumber leaves and resulted in rocaglate resistance in the fungi (Figure 4B). We note that the differential biomass of *C. orbiculare* could not be explained by the damage to cucumber leaves by aglafoline treatment, as no morphological alteration of the leaves was observed under our conditions (Figure 4 — figure supplement 1A). These results demonstrated that the Gly substitution found in *Ophiocordyceps* sp. BRM1 eIF4A provides the molecular basis of antirocaglate properties and allows the growth of the parasitic fungus in the presence of rocaglate (Figure 5).

**Figure 4.**
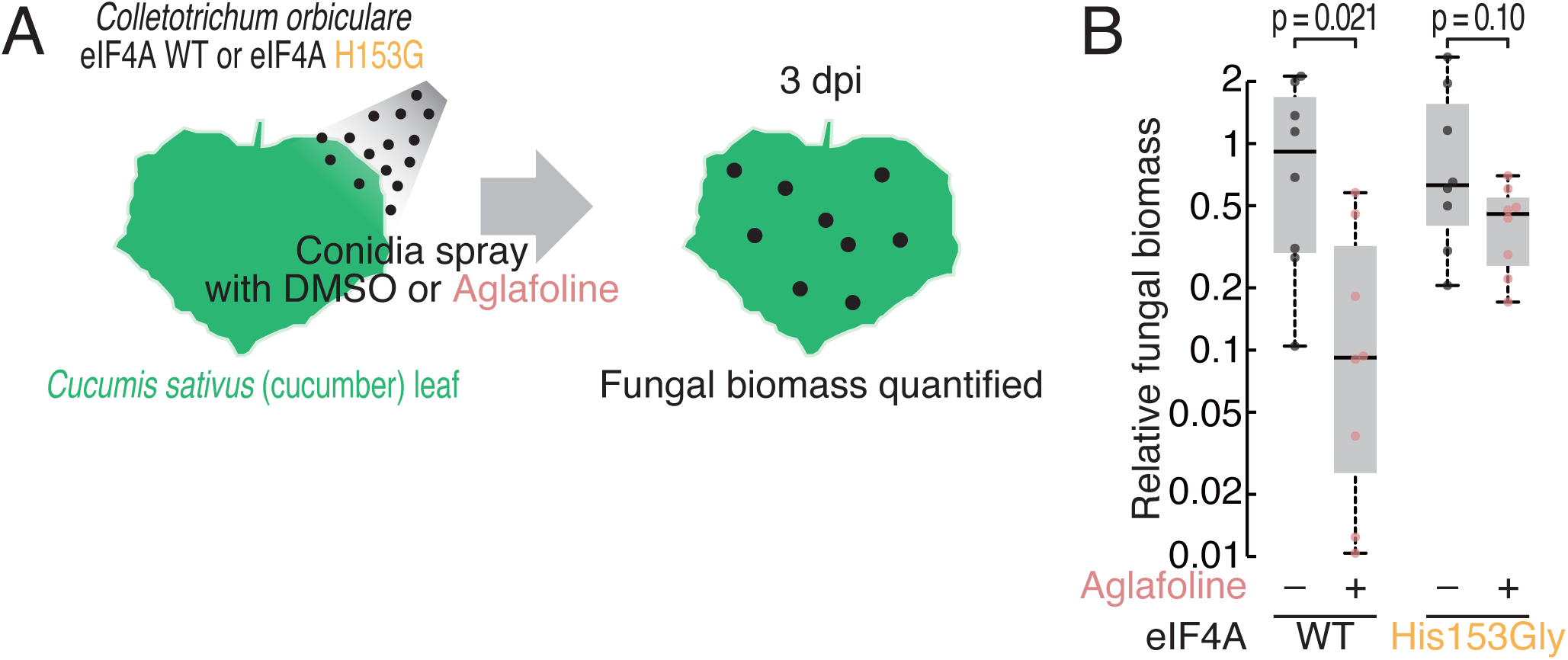
Phenotypic comparison of the *C. orbiculare* eIF4A^WT^ and eIF4A^His153Gly^ strains during infection in the presence of rocaglate. (A) Workflow for monitoring the biomass of *C. orbiculare* eIF4A^WT^ or eIF4A^His153Gly^ strains on cucumber leaves under treatment with aglafoline. (B) Comparison of *in planta* fungal biomass of *C. orbiculare* eIF4A^WT^ or eIF4A^His153Gly^ strains with or without treatment with 1 µM aglafoline. Relative expression levels of the *C. orbiculare 60S ribosomal protein L5* gene (GenBank: Cob_v000942) normalized to that of a cucumber *cyclophilin* gene (GenBank: AY942800.1) were determined by RT– qPCR at 3 dpi (n = 8). The relative fungal biomasses of *C. orbiculare* eIF4A^WT^ and eIF4A^His153Gly^ with aglafoline were normalized to those of eIF4A^WT^ and eIF4A^His153Gly^ without aglafoline, respectively. Significance was calculated by Student’s t test (two-tailed). Three independent experiments showed similar results.

**Figure 5.**
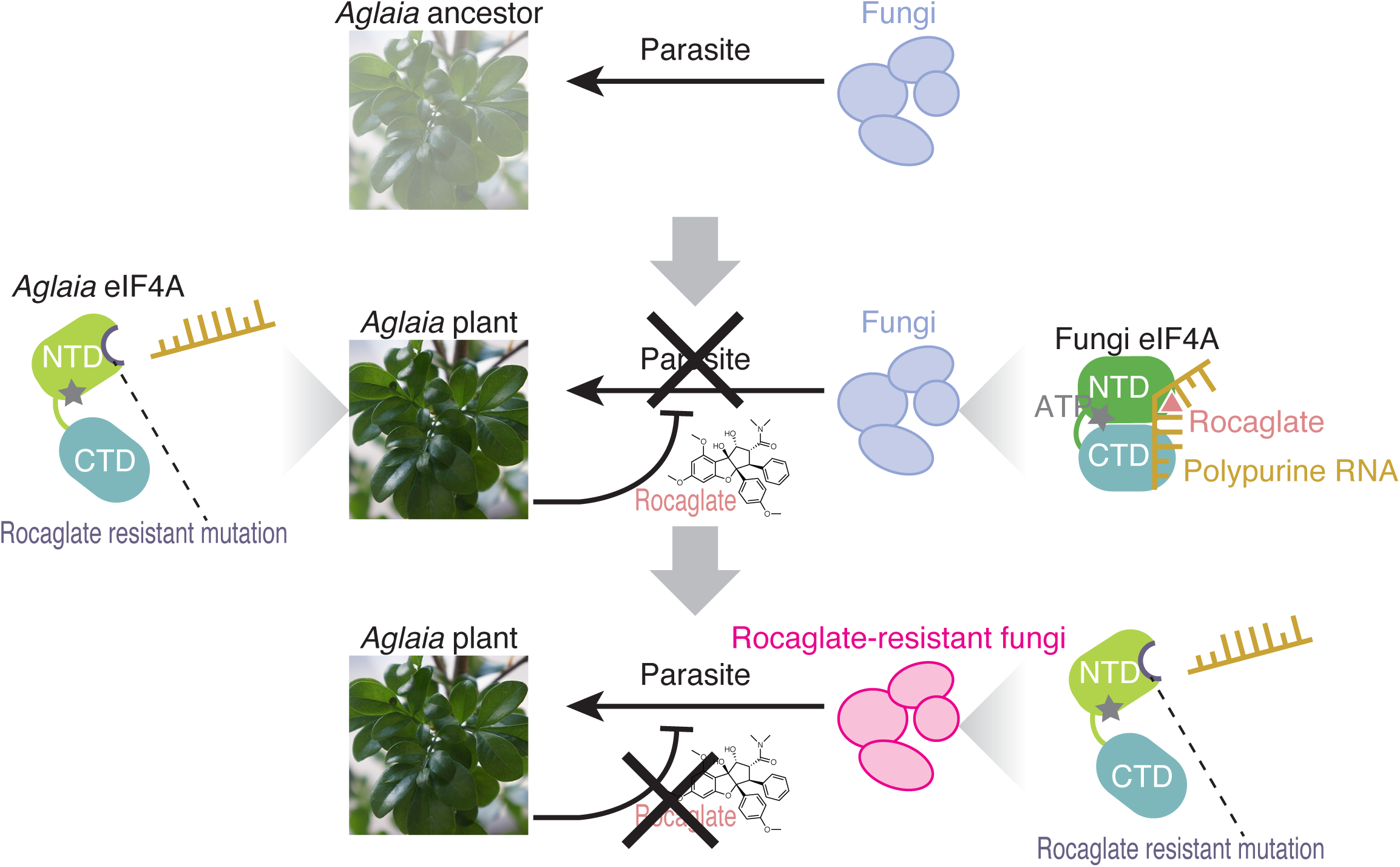
Model of the plant-fungus arms race evoked by rocaglates. The ancestors of the *Aglaia* plants may have been subjected to fungal infection. To counteract this, *Aglaia* plants may have developed rocaglates to target the conserved translation factor eIF4A and to suppress *in planta* fungal growth. Simultaneously, *Aglaia* plants exhibit amino acid substitutions in the rocaglate binding pocket of eIF4As to prevent self-poisoning. Some fungi may impede rocaglate toxin by converting eIF4A to a rocaglate-insensitive form, enabling them to parasitize these plants.

## Discussion

Since plants often produce antifungal secondary metabolites, a specific compound in the host plant may define the interaction between that plants and parasitic fungi (Pusztahelyi et al., 2015). The antifungal activity of rocaglates may protect *Aglaia* plants from phytopathogenic fungi (Figure 5 top and middle). Rocaglate may suppress protein synthesis from survival-essential genes such as those for carbohydrate metabolism and cell wall organization. To survive the presence of rocaglate, which targets the general translation initiation factor eIF4A, this plant converts eIF4A with specific amino acid substitutions (Phe163Leu-Ile199Met: hereafter, we use the human position to specify amino acid residues) to evade the toxicity of the compounds (Iwasaki et al., 2019). This study showed that the parasitic fungus *Ophiocordyceps* sp. BRM1, which possibly originates from *Ophiocordyceps* spp. with endophyte life stage, on *Aglaia* could also overcome this barrier by introducing an amino acid conversion (Phe163Gly) in eIF4A (Figure 5 bottom). Our results highlighted a tug-of-war between host plants and parasitic fungi through the production of translation inhibitory compounds and mutagenization in the target translation factor.

The molecular basis of secondary metabolite resistance in *Ophiocordyceps* sp. BRM1 is markedly distinct from the known strategies developed in other fungi. Avenacin from oats — an example of a plant-secreted antimicrobial substance (Morrissey and Osbourn, 1999) — is a triterpenoid that forms complexes with sterols in fungal cell membranes, causes a loss of membrane integrity, and thus exerts an antifungal effect (Armah et al., 1999; Osbourn et al., 1994). To counteract this compound and infect oats, the phytopathogenic fungus *Gaeumannomyces graminis* var. *avenae* (*Gga*) secretes avenaciase (Crombie et al., 1986; Osbourn et al., 1995), a β-glycosyl hydrolase that hydrolyzes terminal D-glucose in the sugar chain of avenacin. Indeed, avenacin degradation by the enzyme determines the host range of the fungus (Bowyer et al., 1995). In contrast to the detoxification strategy, *Ophiocordyceps* sp. BRM1 may cope with rocaglates through desensitization of the target protein eIF4A with an amino acid substitution (Figure 2), leaving the compound intact.

The different resistance mechanisms to toxic small molecules should be highly related to the compound targets. Since sterols targeted by avenacin are biosynthesized via complicated multiple steps with divergent enzymes and their diverse structures are determined by the requirements of the membrane functions, the conversion of target sterols to evade avenacin requires many enzyme modifications and rarely occurs. On the other hand, the target of rocaglates is an eIF4A protein (and DDX3 protein, see below for details), and thus, evasion by a single amino acid mutation is relatively likely. These results exemplify the mechanistic diversity of attack and counterattack during plant-fungal pathogen interactions.

Although we observed that Gly163 in *Ophiocordyceps* sp. BRM1 eIF4A produced a substantial change in sensitivity to rocaglate, the resistance may not be as complete as that obtained by the substitution found in *Aglaia* Phe163Leu (Iwasaki et al., 2019). Additionally, DDX3, which was recently found to be an alternative target of rocaglate (Chen et al., 2021), did not have amino acid replacements in *Ophiocordyceps* sp. BRM1 (Figure 4 — figure supplement 1B), whereas *Aglaia* DDX3s bear replacements at Gln360 (Chen et al., 2021). This may indicate that *Ophiocordyceps* sp. BRM1 is still in the process of evolving fitness for growth in *Aglaia* plants. Alternatively, the rocaglate-resistant amino acid conversions may involve a trade-off with lower basal translation activity, as we observed in the human reconstitution system (Figure 3 — figure supplement 1A). Even with the inefficiency in translation, given that other fungi could not use the resources from the plant, this substitution may still be beneficial to fungi because of the lack of competition from other fungal species. These possibilities are not mutually exclusive.

## Acknowledgments

We thank all the members of the Iwasaki laboratory for constructive discussions, technical help, and critical reading of the manuscript. We are also grateful to the Support Unit for Bio-Material Analysis, RIKEN CBS Research Resources Division for technical help, supercomputer HOKUSAI Sailing Ship in RIKEN for computation, and the Vincent J. Coates Genomics Sequencing Laboratory (supported by National Institutes for Health Instrumentation Grant, S10 OD018174) at UC Berkeley for DNA sequencing. S.I. was supported by the Japan Society for the Promotion of Science (JSPS) (a Grant-in-Aid for Scientific Research [B] JP19H02959), the Ministry of Education, Culture, Sports, Science and Technology (MEXT) (a Grant-in-Aid for Transformative Research Areas [B] “Parametric Translation”, JP20H05784), the Japan Agency for Medical Research and Development (AMED) (AMED-CREST, JP21gm1410001), and RIKEN (“Biology of Intracellular Environments” and “Integrated life science research to challenge super aging society”). T.I. was supported by JSPS (a Grant-in-Aid for Scientific Research [B], JP19H03172), AMED (AMED-CREST, JP21gm1410001), and RIKEN (“Dynamic Structural Biology”, “Biology of Intracellular Environments”, and “Integrated life science research to challenge super aging society”). K.S. was supported by JSPS (a Grant-in-Aid for Scientific Research [S], JP17H06172). N.K. was supported by JSPS (a Grant-in-Aid for Young Scientists, JP20K15500 and JP18K14440), the Japan Science and Technology Agency (JST) (ACT-X, JPMJAX20B4), and RIKEN (Special Postdoctoral Researchers and Incentive Research Projects). Y.S. was supported by JSPS (a Grant-in-Aid for Early-Career Scientists, JP21K15023), MEXT (a Grant-in-Aid for Transformative Research Areas [A] “Multifaceted Proteins”, JP21H05734), and RIKEN (Special Postdoctoral Researchers and Incentive Research Projects). M.C. was an International Program Associate of RIKEN. N.K. and Y.S. were recipients of the Special Postdoctoral Researchers Program of RIKEN.

## Author contributions

Conceptualization, M.C., N.K., and S.I.;

Methodology, M.C., N.K., R.M., Y.S., M.N., M.M., P.G., K.S., T.I., and S.I.;

Formal analysis, M.C., N.K., R.M., Y.S., M.N., M.M., P.G., T.I., and S.I.;

Investigation, M.C., N.K., R.M., Y.S., M.N., M.M., P.G., T.I., and S.I.;

Resources, R.M., P.G., and N.T.I.;

Writing – Original Draft, M.C., N.K., and S.I.;

Writing – Review & Editing, M.C., N.K., R.M., Y.S., M.N., M.M., P.G., N.T.I., K.S., T.I., and S.I.;

Visualization, M.C., N.K., Y.S., T.I., and S.I.;

Supervision, Y.S., N.T.I., K.S., T.I., and S.I.;

Project administration, S.I.;

Funding Acquisition, N.K., Y.S., K.S., T.I., and S.I.

## Materials and Methods

### RNA-Seq and *de novo* transcriptome assembly of *Ophiocordyceps* sp. BRM1

Fungi on the stem of *Aglaia odorata* (grown in Berkeley, California) were harvested and subjected to RNA extraction with hot phenol. After further chloroform extraction, RNA was subjected to rRNA depletion by a Ribo-Zero Gold rRNA Removal Kit (Yeast) (Illumina). The RNA-Seq library was generated by a TruSeq Stranded mRNA Kit (Illumina) and sequenced by HiSeq4000 (Illumina) with a paired-end 100-bp option. Notably, reads from rRNA genes [*i.e.*, internal transcribed spacer (ITS)] remained after rRNA depletion and were used for phylogenetic analysis.

Transcriptome assembly and functional annotation were performed as described previously (Iwasaki et al., 2019) using Trinity (Grabherr et al., 2011) and Trinotate (Haas et al., 2013). The eIF4A and DDX3 homologous sequences were aligned with MUSCLE (https://www.ebi.ac.uk/Tools/msa/muscle/) and depicted by ESPript 3.0 (http://espript.ibcp.fr/ESPript/ESPript/). eIF4A and DDX3 homologous sequences of model species were obtained from UniProt. For *Ophiocordyceps* species, *Tolypocladium* species, and *Colletotrichum orbiculare*, the ORF databases were obtained from EnsemblFungi (https://fungi.ensembl.org/index.html) or the Ohm laboratory (http://fungalgenomics.science.uu.nl) (de Bekker et al., 2017). The closest homologs registered in UniProt were searched by BLASTp (Camacho et al., 2009) (https://ftp.ncbi.nlm.nih.gov/blast/executables/blast+/LATEST/) for all the proteins in the databases to survey the eIF4A and DDX3 homologs in each species.

### Phylogenetic analysis

To identify the genus of the *Aglaia*-infecting fungus, closely related species were predicted. The *de novo*-assembled transcriptome sequence of the *Aglaia*-infecting fungus was searched by BLASTn (Camacho et al., 2009) (https://ftp.ncbi.nlm.nih.gov/blast/executables/blast+/LATEST/) using the *Colletotrichum aotearoa* ICMP 18537 ITS sequence (GenBank accession: NR_120136) (Schoch et al., 2014) as a query. Using the best hit sequence as a query, BLASTn was performed against the NCBI nucleotide collection (nr/nt) (Supplementary Table 2).

A multilocus phylogenetic analysis of the *Aglaia*-infecting fungus with *Ophiocordyceps* species was performed. A total of 68 isolates were used for phylogenetic analysis, including an *Aglaia*-infecting fungus, 63 previously classified *Ophiocordyceps* strains consisting of 52 species, and 4 *Tolypocladium* species that were expected to serve as outgroups (Supplementary Table 3). DNA sequences of ITS, SSU, LSU, *TEF1a*, and *RPB1* were used as previously reported for the classification of *Ophiocordyceps* species (Xiao et al., 2017). Additional genomic sequences of *Ophiocordyceps* species identified by BLASTn were added to the analysis (Supplementary Table 3). A phylogenetic tree was calculated following previously described methods (Gan et al., 2017; Xiao et al., 2017). Each locus (ITS, LSU, SSU, *RPB1*, and *TEF1α*) of 68 isolates (Ban et al., 2015; Castlebury et al., 2004; Chen et al., 2013; de Bekker et al., 2017; Hu et al., 2013; Kepler et al., 2012; Liu et al., 2002; Luangsa-Ard et al., 2011, 2010; Quandt et al., 2018, 2014; Sanjuan et al., 2015; Schoch et al., 2012; Spatafora et al., 2007; Sung et al., 2007; Wen et al., 2013; Will et al., 2020; Xiao et al., 2017) was aligned using MAFFT v7.480 (Katoh and Standley, 2013) and trimmed by trimAl v1.4.rev15 (Capella-Gutiérrez et al., 2009) with an automated setting. The processed sequences obtained from every 68 isolates were concatenated by catfasta2phyml (https://github.com/nylander/catfasta2phyml) to generate sequences comprising 3,910 nucleotide positions, including gaps (gene boundaries ITS, 1-463; LSU, 464-1,363; SSU, 1,364-2,248; *RPB1*, 2,249-2,922; *TEF1α*, 2,923-3,910). The best model for nucleotide substitutions under the BIC criterion was determined by ModelTest-NG v.0.1.6 (https://github.com/ddarriba/modeltest) as follows: ITS, TIM3ef+G4; LSU, TIM1+I+G4; SSU, TPM3+I+G4; *RPB1*, TIM1+I+G4; and *TEF1α*, TrN+I+G4. Then, the maximum-likelihood phylogeny was estimated based on concatenated sequences by RAxML-NG v.0.9.0 (https://github.com/amkozlov/raxml-ng) using the ModelTest-NG specified best models for each partition with 1,000 bootstrap replicates. The best-scoring maximum-likelihood trees with bootstrap support values were visualized in iTOL v6 (https://itol.embl.de/). Given sufficient separation from other known *Ophiocordyceps*, the fungus was named *Ophiocordyceps* sp. BRM1.

### Compounds

RocA (Sigma–Aldrich) and aglafoline (MedChemExpress) were dissolved in dimethyl sulfoxide (DMSO) and used for this study.

### Plasmid construction

#### pColdI-H. sapiens eIF4A1 WT, Phe163Gly, and Phe163His

pColdI-*H. sapiens* eIF4A1 has been reported previously (Iwasaki et al., 2019). Phe163Gly and Phe163His substitutions were induced by site-directed mutagenesis.

#### pColdI-O. sinensis eIF4A1 WT and His154Gly

DNA fragments containing the *O. sinensis* eIF4A1 gene were synthesized by Integrated DNA Technologies (IDT) and inserted into pColdI (TaKaRa) downstream of the His tag with In-Fusion HD (TaKaRa). The His154Gly substitution was induced by site-directed mutagenesis.

#### pColdI-Ophiocordyceps sp. BRM1 eIF4A iso4 WT and Gly172Phe

The cDNA library of *Ophiocordyceps* sp. BRM1 was reverse-transcribed with ProtoScript II Reverse Transcriptase (New England Biolabs) and Random Primer (nonadeoxyribonucleotide mix: pd(N)_9_) (TaKaRa) from the total RNA of *Ophiocordyceps* sp. BRM1 (see details in the section “RNA-Seq and *de novo* transcriptome assembly for *Ophiocordyceps* sp. BRM1”). Using the cDNA as a template, DNA fragments containing eIF4A iso4 were PCR-amplified and inserted into pColdI (TaKaRa) downstream of the His tag with In-Fusion HD (TaKaRa). The Gly172Phe substitution was induced by site-directed mutagenesis.

#### pENTR4-C. orbiculare eIF4A WT and His153Gly

To replace the *eIF4A* (GenBank: Cob_v000942) sequence in the *C. orbiculare* genome with synthesized *C. orbiculare eIF4A* WT or His153Gly, donor DNAs for homology-directed repair were constructed. DNA fragments, including 2-kb genome sequences upstream and downstream of *C. orbiculare eIF4A* (as homology arms), the *C. orbiculare eIF4A* genome sequence, and the neomycin phosphotransferase II (*NPTII*) expression cassette, were fused into the pENTR4 plasmid (Thermo Fisher Scientific) by HiFi DNA assembly (New England Biolabs). These fragments were PCR-amplified using *C. orbiculare* genomic DNA, which was isolated from the mycelium, or pII99 plasmid (Namiki et al., 2001). The His153Gly substitution was induced by site-directed mutagenesis. sgRNA-targeted sequences in homology arm sequences were deleted by site-directed deletion to prevent cleavage by CRISPR–Cas9.

### Recombinant protein purification

His-tagged recombinant proteins were purified as described previously (Chen et al., 2021). BL21 Star (DE3) (Thermo Fisher Scientific) cells were transformed with pColdI plasmids (see “Plasmid construction” section). After the induction of protein expression by isopropyl-*β*-D-thiogalactopyranoside (IPTG) at 15°C overnight, cells were collected by centrifugation and flash-frozen in liquid nitrogen. Subsequently, the thawed cells were lysed by sonication.

The His-tagged protein was purified by Ni-NTA agarose (Qiagen). Eluted proteins from beads were then applied to the NGC chromatography system (Bio–Rad). Using a HiTrap Heparin HP column (1 ml, GE Healthcare), proteins were fractionated via an increased gradient of NaCl. The peak fractions were collected, buffer-exchanged by NAP-5 or PD-10 (GE Healthcare) into the storage buffer [20 mM HEPES-NaOH pH 7.5, 150 mM NaCl, 10% glycerol, and 1 mM dithiothreitol (DTT)], concentrated with a Vivaspin 6 (10 kDa MWCO) (Sartorius), flash-frozen by liquid nitrogen, and then stored at −80°C.

### Fluorescence polarization assay

Fluorescence polarization assay was performed as previously described (Chen et al., 2021). The reaction was prepared with 0-25 μM recombinant protein, 10 nM FAM-labeled [AG]_10_ RNA, 1 mM AMP-PNP, 1 mM MgCl_2_, 20 mM HEPES-NaOH (pH 7.5), 150 mM NaCl, 1 mM DTT, 5% glycerol, and 1% DMSO (as a solvent of RocA) with or without 50 μM RocA/aglafoline. After incubation at room temperature for 30 min, the mixture was transferred to a black 384-well microplate (Corning), and the fluorescence polarization was measured by an Infinite F-200 PRO (Tecan). Under ADP + Pi conditions, 1 mM ADP and 1 mM Na_2_HPO_4_ were used as substitutes for AMP-PNP. The data were fitted to the Hill equation to calculate *K_d_* and visualized by Igor Pro 8 (WaveMetrics). The affinity fold change was calculated as the fold reduction of *K_d_* in RocA compared to *K_d_* in DMSO.

### *In vitro* translation assay in reconstituted system

The reconstitution system for human translation has been described previously (Iwasaki et al., 2019; Machida et al., 2018; Yokoyama et al., 2019). The DNA fragments PCR-amplified from psiCHECK2-7×AGAGAG motifs were used as a template for *in vitro* transcription with a T7-Scribe Standard RNA IVT Kit (CELLSCRIPT). Following capping and poly(A)-tailing with the ScriptCap m^7^G Capping System, a ScriptCap 2′-*O*-Methyltransferase Kit, and an A-Plus Poly(A) Polymerase Tailing Kit (CELLSCRIPT), mRNA was used for the translation assay. The *in vitro* translation reaction and luciferase assay were performed as previously described (Iwasaki et al., 2019) with some modifications. The final concentrations of mRNA and the eIF4A protein were 100 ng/µl and 2.45 µM, respectively. The translation mixture was incubated for 2.5 h. The fluorescence signal was detected using the Renilla-Glo Luciferase Assay System (Promega) and counted in an EnVision 2104 plate reader (PerkinElmer).

### Fungal transformation

*C. orbiculare* strain 104-T (NARO Genebank ID: MAFF 240422), a causal agent of anthracnose disease in Cucurbitaceae plants, was used. The isolated strains in this study are also listed in Supplementary Table 4.

#### Preparation of protoplasts

*C. orbiculare* protoplasts were prepared as previously described (Kubo, 1991; Rodriguez and Yoder, 1987; Vollmer and Yanofsky, 1986) with modifications. A frozen glycerol stock of *C. orbiculare* was streaked on 3.9% (w/v) potato dextrose agar (PDA) medium (Nissui) in a 90-mm dish and incubated at 25°C in the dark for 3 d. Outer edges of a colony were transferred to 20 ml of 2.4% (w/v) potato dextrose broth (BD) and incubated for 2 d at 25°C in the dark. The proliferated mycelium was collected using a 70-µm cell strainer (Corning) and incubated in 150 ml of potato-sucrose liquid medium supplemented with 0.2% yeast extract (BD Biosciences) at 25°C with shaking at 140 rpm. The mycelium was harvested, washed with sterile water, and resuspended in 20 ml of filter-sterilized (0.2-µm pore size, GE Healthcare) osmotic medium (1.2 M MgSO_4_ and 5 mM Na_2_HPO_4_) containing 10 mg/ml driselase from *Basidiomycetes* sp. (MERK) and 10 mg/ml lysing enzyme from *Trichoderma harzianum* (MERK) in a 50-ml tube (Falcon, Corning). The suspension was gently agitated in a rotary shaker at 60 rpm for 90 min at 30°C. Then, the suspension was underlaid with 20 ml of trapping buffer (0.6 M sorbitol, 50 mM Tris-HCl pH 8.0, and 50 mM CaCl_2_) and centrifuged at 760 ×g for 5 min using a swinging-bucket rotor (Hitachi, T4SS31). Protoplasts isolated from the interface of the two layers were pelleted, washed twice using STC (1 M sorbitol, 50 mM Tris-HCl pH 8.0, and 50 mM CaCl_2_), resuspended in STC at 10^8^-10^9^ protoplasts/ml, added to a 25% volume of polyethylene glycol (PEG) solution (40% [w/w] PEG3350, 500 mM KCl, 40 mM Tris-HCl pH 8.0, and 50 mM CaCl_2_), and stored at −80°C until use.

#### gRNA preparation

Template DNA fragments for sgRNA *in vitro* transcription were PCR-amplified using the primers listed in Supplementary Table 5. Using the DNA fragments, sgRNAs (sgRNAUP-1, sgRNAUP-2, sgRNADW-1, and sgRNADW-2) were prepared with a CUGA7 gRNA Synthesis Kit (Nippon Gene) following the manufacturer’s protocol.

#### Transformation

The transformation was performed as previously described (Foster et al., 2018; Kubo, 1991; Yelton et al., 1984) with modifications. The mixture of plasmid DNA (5 µg, pENTR4-*C. orbiculare* eIF4A WT or His153Gly), the four sgRNAs (250 ng each), and Cas9 nuclease protein NLS (15 µg, Nippon Gene) were added to 150 µl of *C. orbiculare* protoplasts, followed by the addition of 1 ml of STC and 150 µl of PEG solution. The resulting suspension was incubated for 20 min on ice, supplemented with 500 µl of PEG solution, and gently agitated by hand. The suspension was serially diluted with a second addition of 500 µl, a third addition of 1 ml, and fourth and fifth additions of 2 ml of PEG solution, with gentle agitation at every dilution step. After incubation for 10 min at room temperature, the PEG solution was removed by centrifugation. The protoplasts were resuspended in 1 ml of STC, diluted with 15 ml of regeneration medium (3.12% [w/v] PDA and 0.6 M glucose), and then spread onto a plate containing 40 ml of selection medium (3.9% [w/v] PDA and 0.6 M glucose) containing 200 µg/ml G418 (Fujifilm Wako Chemicals). The plate was incubated for 5 d at 25°C in the dark. The G418-resistant colonies were further seeded in fresh selection medium containing G418 and subjected to selection for an additional 5 d.

#### Screening by PCR

Then, the genomic DNA isolated from each colony was subjected to PCR to ensure the desired transformation (see Figure 3 — figure supplement 1B for the design). The primers used for the PCR screening are listed in Supplementary Table 5. The selected transformant conidia were suspended in 25% glycerol and stored at −80°C until use.

### *Colletotrichum orbiculare* mitochondrial genome assembly

Reads from three PacBio RSII cells of the *C. orbiculare* 104-T whole genome sequencing (Gan et al., 2019) were mapped onto *C. orbiculare* scaffolds that were identified as potential mitochondrial sequences by the NCBI Genomic contamination screen with minimap2 (v2.17-r941) (Li, 2018) using the map-bp setting. Aligned fasta reads were then assembled using flye (2.8.1-b1676) (Kolmogorov et al., 2019) with default settings (min overlap=5000 bp). The assembly (GenBank accession: MZ424187) possessed a 36,318 bp contig with 2023.72× coverage and showed the highest homology to the *Colletotrichum lindemuthianum* completed mitochondrial genome (KF953885) according to nucmer (Delcher et al., 2003). These genome data were used for data processing for ribosome profiling.

### Ribosome profiling

#### Cell culture; mycelia

Glycerol stocks of *C. orbiculare* eIF4A^WT^#1 and eIF4A^H153G^#1 strains were streaked on PDA in 90-mm plastic petri dishes and incubated for 3 d. A single colony of each strain was transferred onto PDA and incubated for 3 d. The outer edges of colonies were transferred to 90-mm plastic dishes filled with 20 ml of PDB using plastic straws and incubated for 4 d. Aglafoline (0.3 or 3 µM) or DMSO was added to dishes and incubated for 6 h.

#### Cell culture; conidia

A single colony from the glycerol stocks was cultured by the same method used for mycelium preparation. The outer edges of colonies of each strain were transferred into six 300-ml flasks filled with 100 ml of PDA. Two milliliters of sterilized water was added to each flask, and the flasks were shaken well to ensure that the mycelial cells adhered to the entire surface of the PDA evenly. After 6 d of incubation in the dark, conidia generated on the surface of PDA were suspended in 20 ml of sterilized water. The conidial suspension was filtered through a 100-µm pore-size cell strainer and collected by centrifugation at 760 ×g for 5 min at room temperature. Twenty milliliters of resuspended conidia at 0.5 OD_600_ (approximately 2.5 × 10^6^ conidia/ml) was dispensed in 50 ml ProteoSave SS tubes (Sumitomo Bakelite Co., Ltd.) and then treated with aglafoline (0.3 or 3 µM) or DMSO for 6 h in the dark with shaking at 140 rpm.

#### Cell harvest

Cells were filtered by an MF membrane (0.45-µm pore size, Millipore), immediately scraped from the filter, and soaked in liquid nitrogen for 30 s. Into a tube holding the cell pellet and liquid nitrogen, 600 µl of lysis buffer (20 mM Tris-HCl pH 7.5, 150 mM NaCl, 5 mM MgCl_2_, 1 mM DTT, 100 µg/ml cycloheximide, 100 µg/ml chloramphenicol, and 1% Triton X-100) was added dropwise to form ice grains. The samples were stored at − 80°C to evaporate the liquid nitrogen.

#### Library preparation

The frozen cells and lysis buffer grains were milled by a Multi-beads Shocker (YASUI KIKAI) at 2,800 rpm for 15 se for 1 cycle. The lysates were thawed on ice and centrifuged at 3000 ×g and 4°C for 5 min. The supernatant was treated with 25 U/ml Turbo DNase (Thermo Fisher Scientific) for 10 min and then clarified by centrifugation at 20,000 ×g and 4°C for 10 min. Then, the supernatant was used for downstream ribosome profiling library preparation as described previously (Mito et al., 2020). Briefly, the lysates containing 10 µg of total RNA were treated with 2 U/µg RNase I (Lucigen) at 25°C for 45 min. After ribosomes were collected by a sucrose cushion, the RNAs were separated in 15% urea PAGE gels, and the RNA fragments ranging from 17 to 34 nt were excised. Subsequently, the RNAs were dephosphorylated and ligated to linkers. Following rRNA removal with a Ribo-Minus Eukaryotes Kit for RNA-Seq (Invitrogen), the RNA fragments were reverse-transcribed, circulated, and PCR-amplified. The final DNA libraries were sequenced on a HiSeq X (Illumina) with a paired-end 150-bp option.

#### Data processing

Sequence data were processed as previously described (McGlincy and Ingolia, 2017) with modifications. Using the fastp (Chen et al., 2018) tool, sequences of reads 1 were corrected by reads 2, and then quality filtering and removing adapter sequences of reads 1 were performed. The adapter-removed reads 1 were splitted by the barcode sequence, mapped to rRNA and tRNA sequences of *C. orbiculare*, which were predicted by RNAmmer (http://www.cbs.dtu.dk/services/RNAmmer/) and tRNA-scan SE (http://lowelab.ucsc.edu/tRNAscan-SE/) in the genome of *C. orbiculare* 104-T (Gan et al., 2019) (PRJNA171217), using STAR 2.7.0a (Dobin et al., 2013), and were removed from analysis. For all predicted tRNAs, the CCA sequence was added to the 3′ end. The remaining reads were mapped to the *C. orbiculare* genome (Gan et al., 2019) by STAR 2.7.0a. The A-site offset of footprints was empirically estimated to be 15 for the 19-21 nt and 24-30 nt footprints. Footprints located on the first and last 5 codons of each ORF were omitted from the analysis.

The translation change induced by aglafoline was calculated by DESeq2 (Love et al., 2014) and renormalized to mitochondrial footprints (as an internal spike-in standard) (Iwasaki et al., 2016).

For GO analysis, IDs of sensitive mRNAs in *C. orbiculare* conidia were converted to IDs of *Saccharomyces cerevisiae* homologs predicted using BLASTp (https://ftp.ncbi.nlm.nih.gov/blast/executables/blast+/LATEST/) (Camacho et al., 2009) and the S288C reference from the *Saccharomyces* Genome Database (SGD). A functional annotation chart for this list was obtained from DAVID (https://david.ncifcrf.gov/home.jsp) (Huang et al., 2009a, 2009b). GO terms with FDR < 0.05 were considered.

For 5′ UTR assignment of *C. orbiculare*, published RNA-Seq data (GSE178879) (Zhang et al., 2021) were aligned to *C. orbiculare* genome by STAR 2.7.0a and then assembled into transcript isoforms by StringTie 2.2.1 (Kovaka et al., 2019). The extensions upstream from annotated start codons were assigned as 5′ UTR. The 5′ UTR of transcripts expressed in conidia and mycelia were obtained individually. For the analysis of polypurine sequence in Figure 3G and Figure 3 — figure supplement 3E, we used the 5′ UTR with highest coverage in StringTie when multiple 5′ UTR isoforms were assigned.

### Fungal inoculation

Fungal inoculation was performed as previously described (Hiruma and Saijo, 2016; Kumakura et al., 2019) with modifications. Cucumber cotyledons were used for *C. orbiculare* inoculation. Seeds of cucumber, *Cucumis sativus* Suyo strain (Sakata Seed Corp.), were planted on a mix of equal amounts of vermiculite (VS Kakou) and Supermix A (Sakata Seed Corp.). Cucumbers were grown at 24°C under a 10 h light/14 h dark cycle using biotrons (NK Systems). Cotyledons were detached from seedlings of cucumbers and inoculated with *C. orbiculare* at 13 d post-germination. *C. orbiculare* strains (eIF4A^WT^#1 and eIF4A^H153G^#1, Supplementary Table 4) were cultured on 100 ml of 3.9% PDA in a 300-ml flask at 25°C for 6 d in the dark. Conidia that appeared on the surface of PDA were suspended in 20 ml of sterilized water, filtered through cell strainers (100-µm pore size, Corning), pelleted by centrifugation at 760 ×g for 5 min, and resuspended in sterilized water. The concentration of conidia was measured with disposable hemacytometers (Funakoshi) and adjusted to 10^5^ conidia/ml with or without aglafoline (1 µM). Both conidial suspensions contained DMSO at 0.005% (v/v). Conidial suspensions were sprayed onto detached cotyledons using a glass spray (Sansho) and an air compressor (NRK Japan). Inoculated leaves were placed in plastic trays and incubated at 100% humidity for 3.5 d under the same conditions used for plant growth. Using a 6-mm trepan (Kai Medical), 6 leaf discs (LDs) were cut from each leaf, and 48 LDs were collected per sample. Six LDs were placed in a 2-ml steel top tube (BMS) with a Φ5-mm zirconia bead (Nikkato), and eight tubes were prepared for each sample (n = 8). Samples were frozen in liquid nitrogen, ground at 1,500 rpm for 2 min using Shakemaster NEO (BMS), and stored at −80°C until RNA extraction.

### Quantification of fungal biomass *in planta*

The living fungal biomass in cucumber leaves at 3.5 d postinoculation (dpi) was measured by RT–qPCR. Relative expression levels of the *C. orbiculare 60S ribosomal protein L5* gene (GenBank: Cob_v000942) (Gan et al., 2013) normalized to that of a cucumber *cyclophilin* gene (GenBank: AY942800.1) (Liang et al., 2018) were determined. Total RNA was extracted with the Maxwell RSC Plant RNA Kit (Promega) and Maxwell RSC 48 Instrument (Promega) with the removal of genomic DNA according to the manufacturer’s protocol. cDNA was synthesized from 500-1,000 ng of total RNA per sample with a ReverTraAce qPCR RT Kit (TOYOBO) following the manufacturer’s instructions. All RT-qPCRs were performed with THUNDERBIRD Next SYBR qPCR Mix (TOYOBO) and an MX3000P Real-Time qPCR System (Stratagene). The primers used are listed in Supplementary Table 5.

### Resource availability

The results of ribosome profiling (GEO: GSE200060) for *C. orbiculare* and RNA-Seq for *Ophiocordyceps* sp. BRM1 (SRA: PRJNA821935) obtained in this study have been deposited in the National Center for Biotechnology Information (NCBI) database. The *C. orbiculare* mitochondrial genome assembly generated in this study was deposited under accession number MZ424187. The scripts for deep sequencing data analysis were deposited in Zenodo (DOI: 10.5281/zenodo.6787991). Further information and requests for resources and reagents should be directed to and will be fulfilled by the Lead Contact, Shintaro Iwasaki.

---

**Figure 1 — figure supplement 1.**
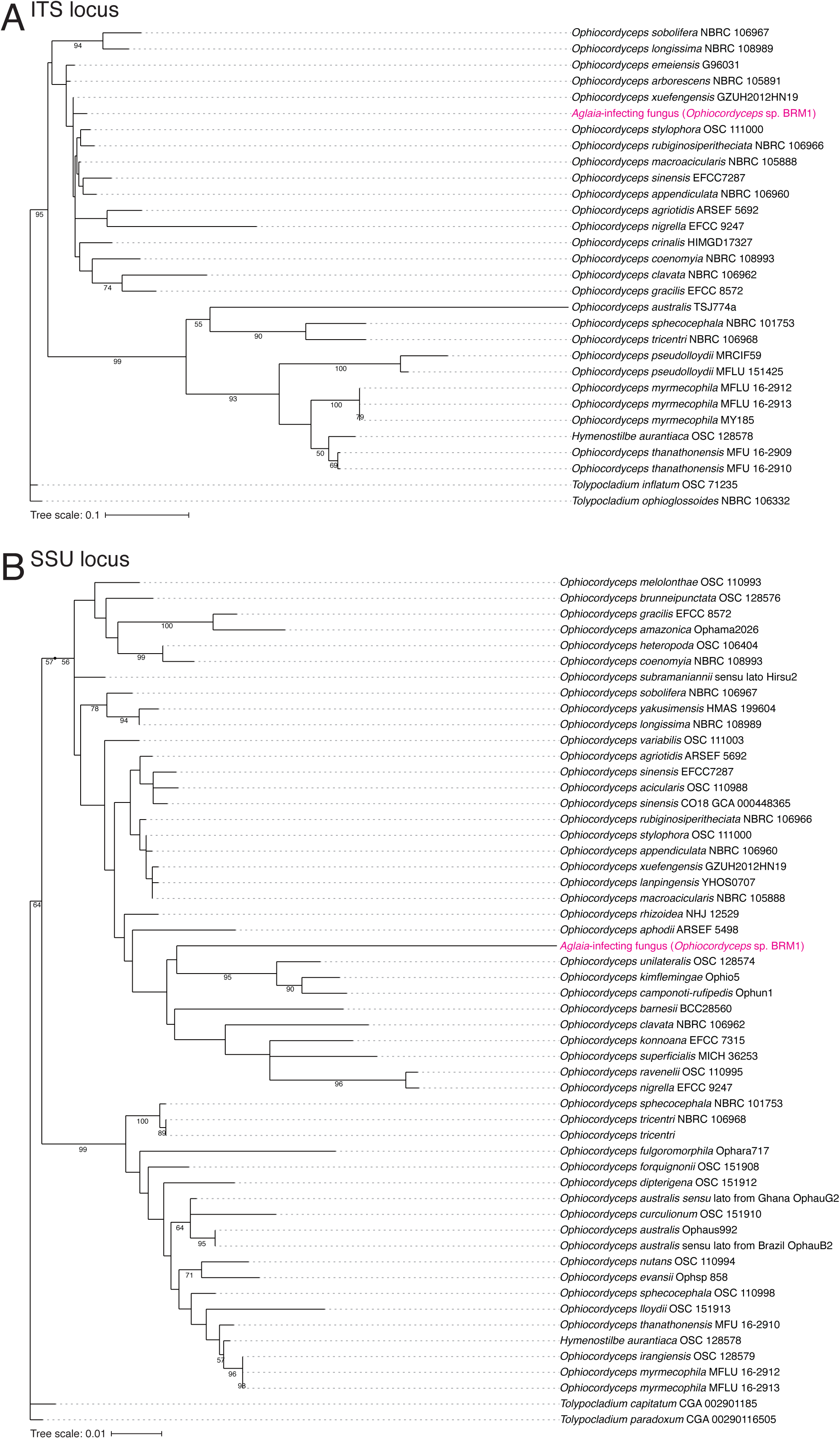
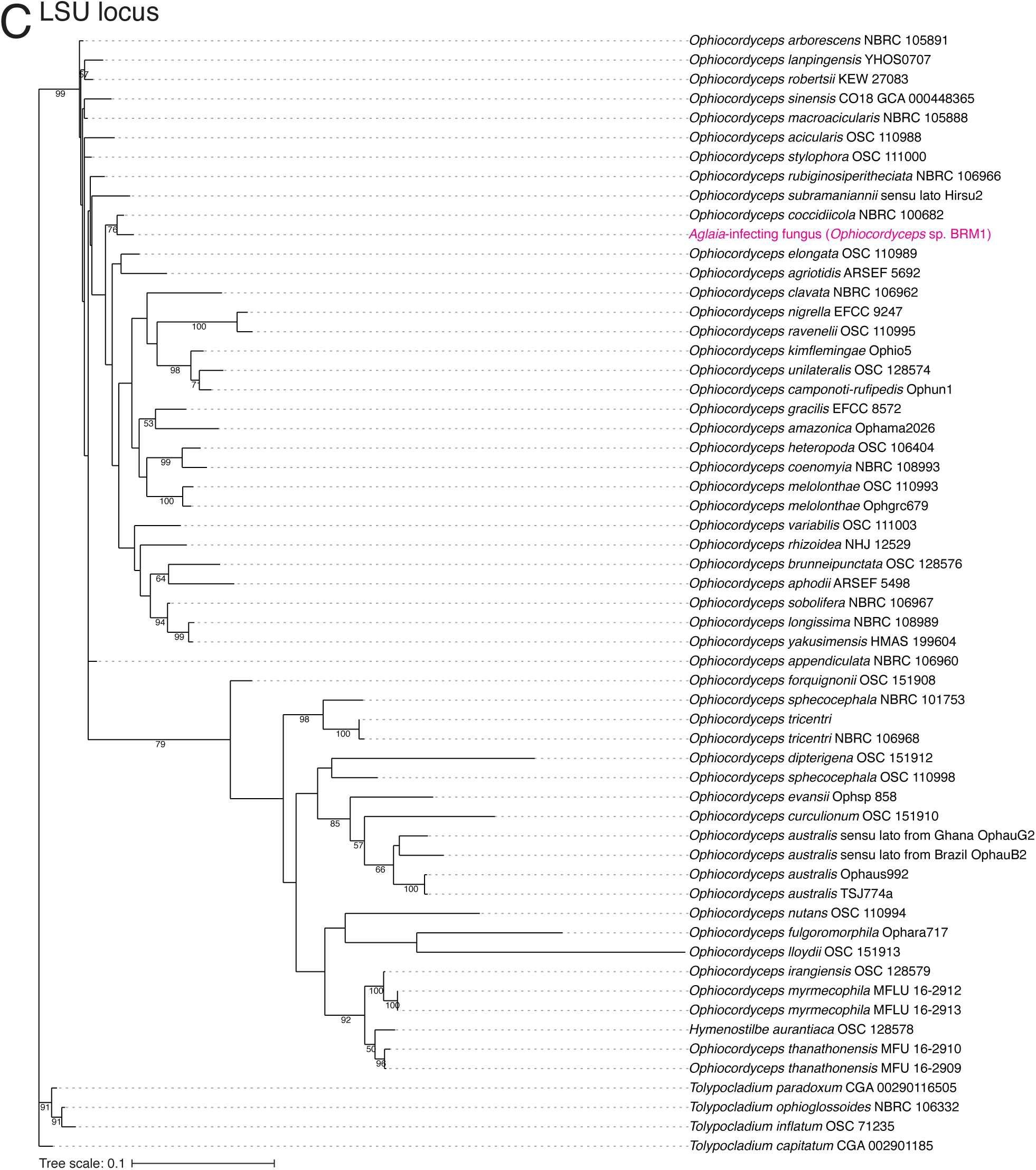
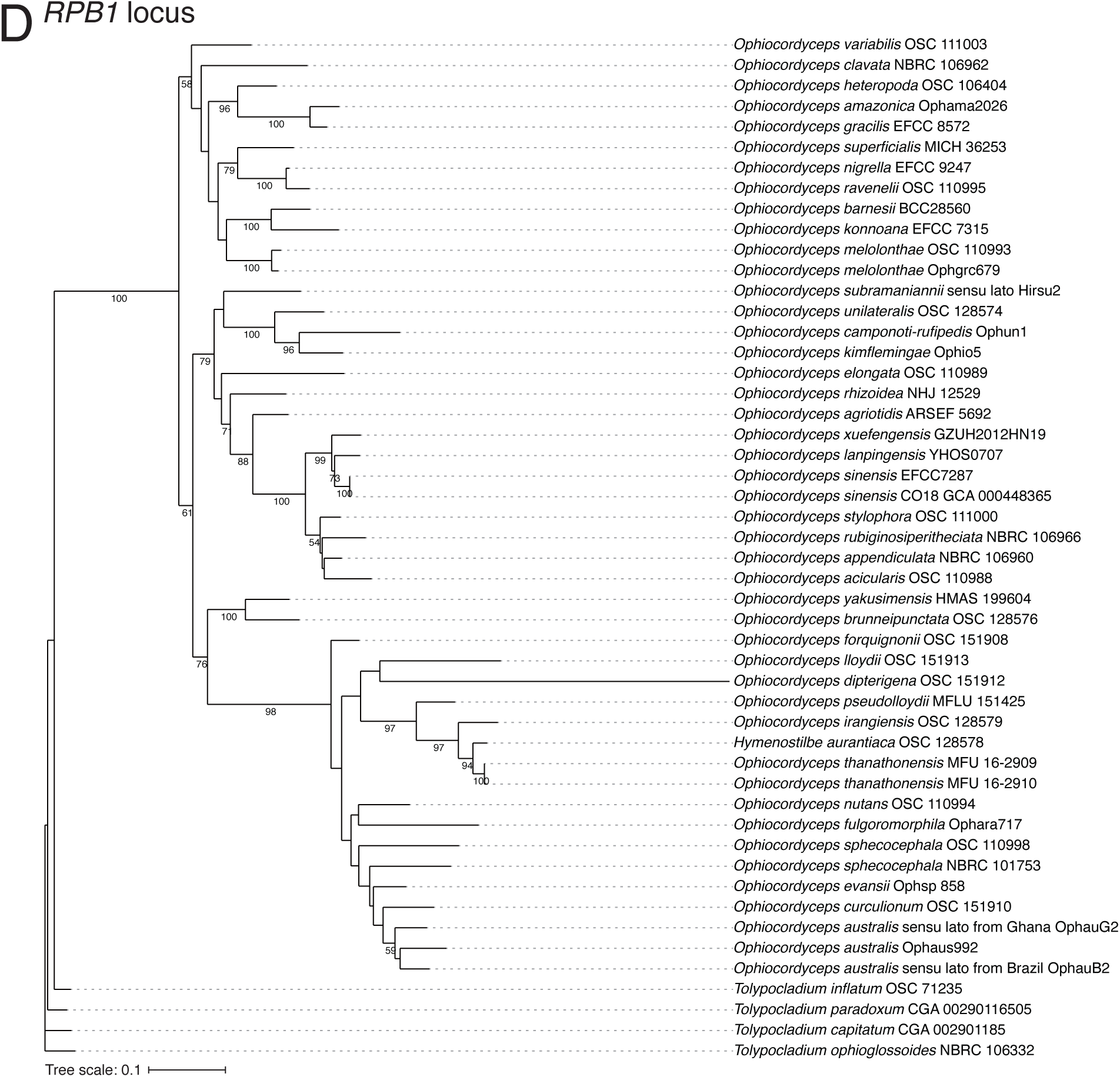
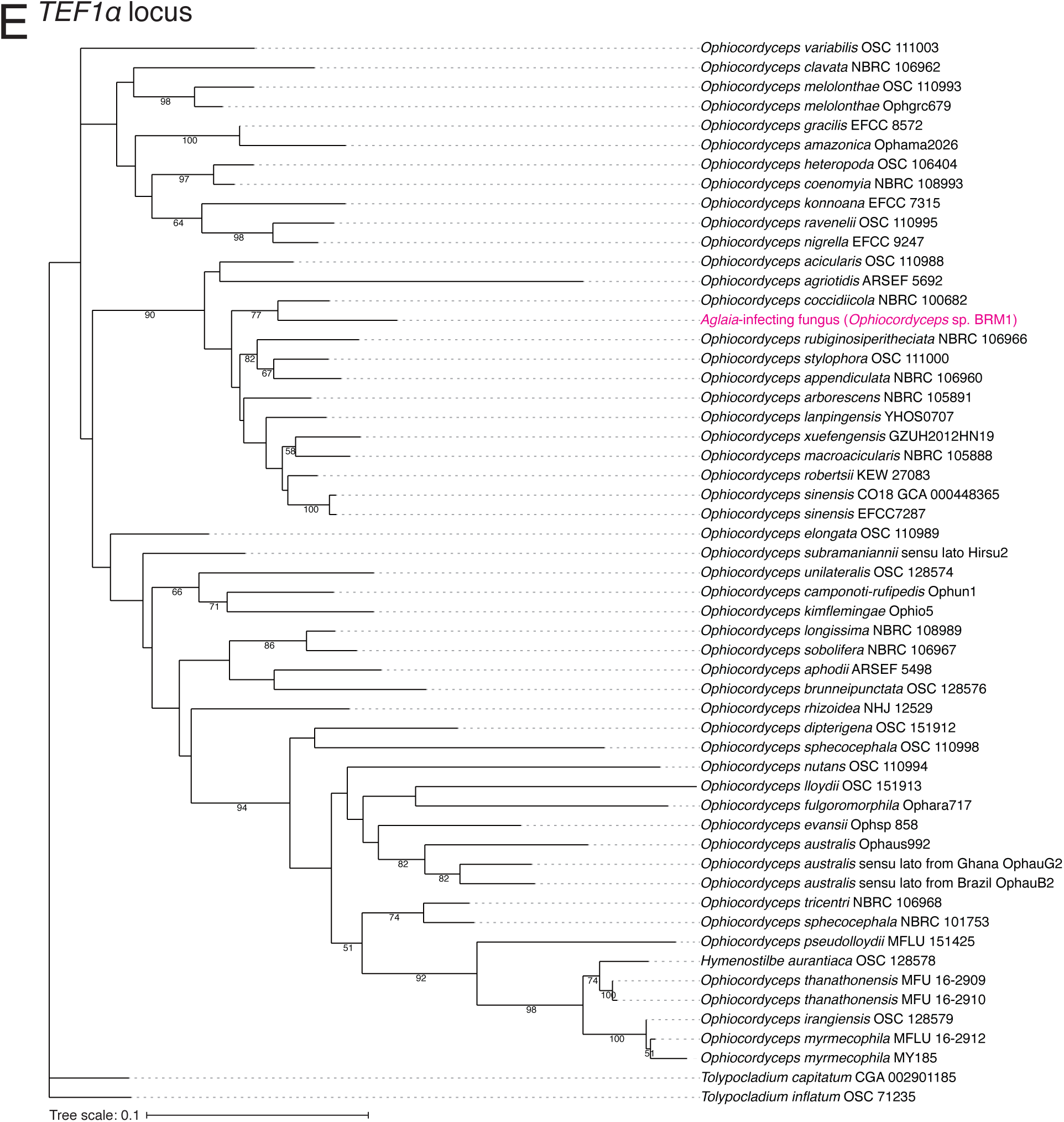
Assessment of *Aglaia*-infecting fungus species. (A-E) The maximum likelihood best scoring trees based on the indicated single gene locus from the *Ophiocordyceps* species with *Tolypocladium* species as outgroups. Numbers at nodes are percentages of bootstrap support values out of 1,000. Only bootstrap values above 50% are shown.

**Figure 2 — figure supplement 1.**
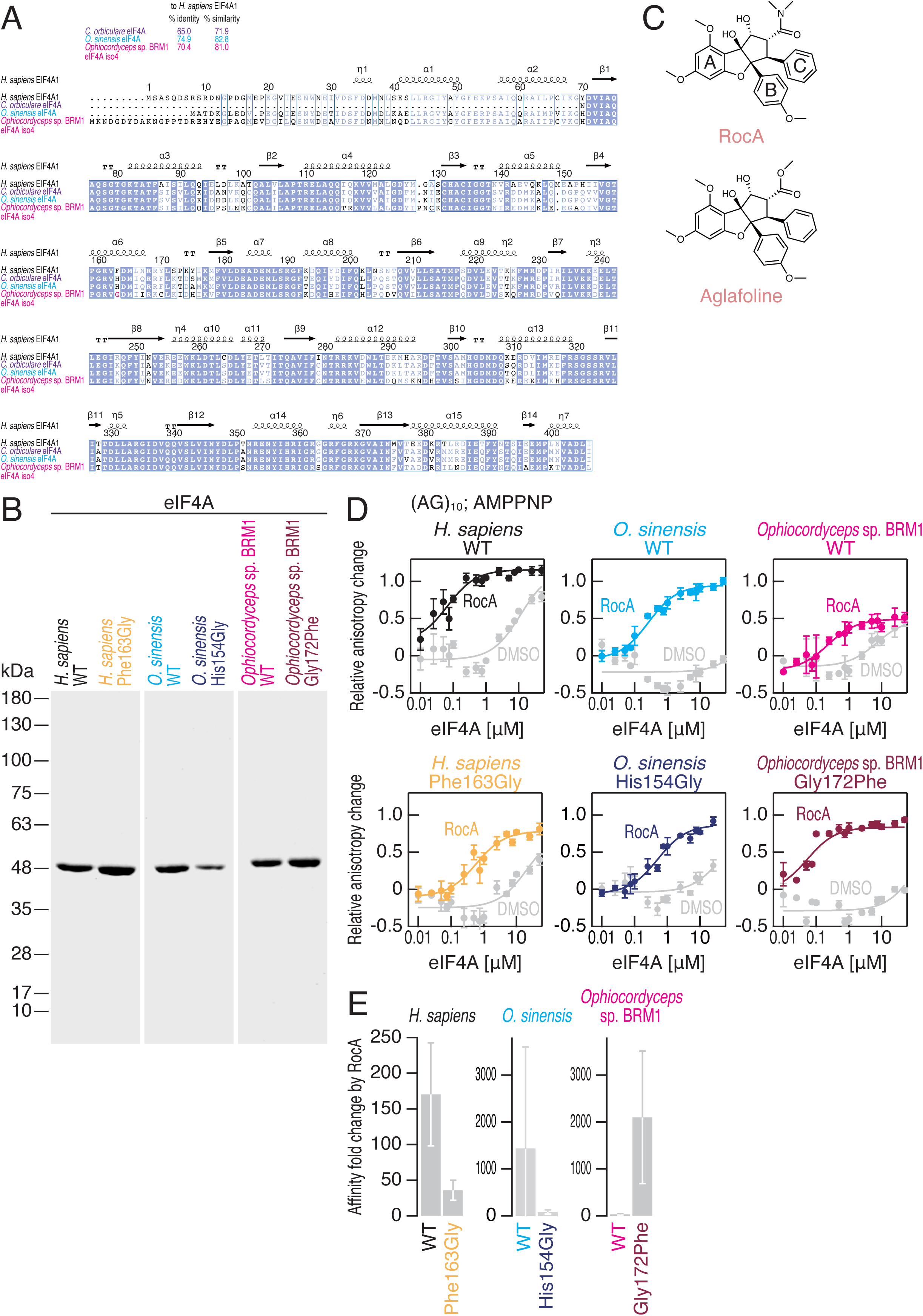
Characterization of recombinant proteins used in this study. (A) Alignments of eIF4A protein sequences for indicated species. Percent similarity and percent identity to *H. sapiens* eIF4A eIF4A1 are shown on the top. (B) Coomassie brilliant blue staining of recombinant eIF4A proteins used in this study. (C) Chemical structures of rocaglates used in this study. (D) Fluorescence polarization assay for FAM-labeled RNA ([AG]_10_) (10 nM). WT and mutated eIF4A proteins from the indicated species were used. An ATP ground state analog AMP-PNP (1 mM) was included in the reaction with RocA (50 µM). Data represent the mean and s.d. (n = 3). (E) Affinity fold changes by amino acid substitutions in C were calculated. Data represent the mean and s.d.

**Figure 3 — figure supplement 1.**
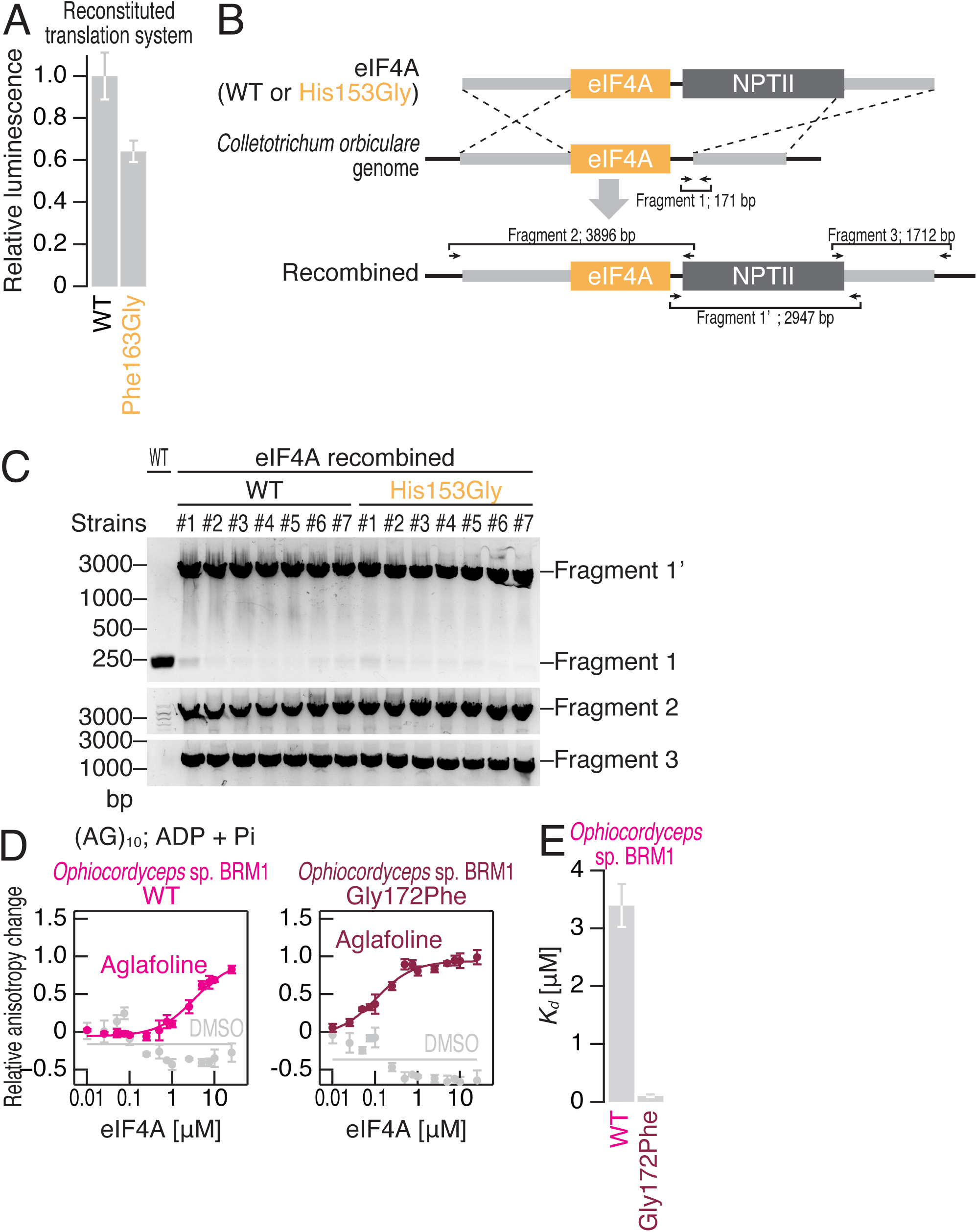
Establishment of eIF4A-engineered *C. orbiculare* strains. (A) Basal protein synthesis activity in an *in vitro* translation system with human factors. Recombinant proteins of *H. sapiens* eIF4A1 WT or Phe163Gly were added to the reaction. (B) Schematics of eIF4A recombination in *C. orbiculare*. *NPTII*, neomycin phosphotransferase II. (C) PCR-based screening of the recombined strains. The primer sets used for screening are depicted in B. (D) Fluorescence polarization assay for FAM-labeled RNA ([AG]_10_) (10 nM). WT and mutated *Ophiocordyceps* sp. BRM1 eIF4A proteins from the indicated species were used. To measure ATP-independent RNA clamping induced by aglafoline (50 µM), ADP and Pi were included in the reaction. Data represent the mean and s.d. (n = 3). (E) The summary of *K_d_* in D under aglafoline is depicted. Data represent the mean and s.d.

**Figure 3 — figure supplement 2.**
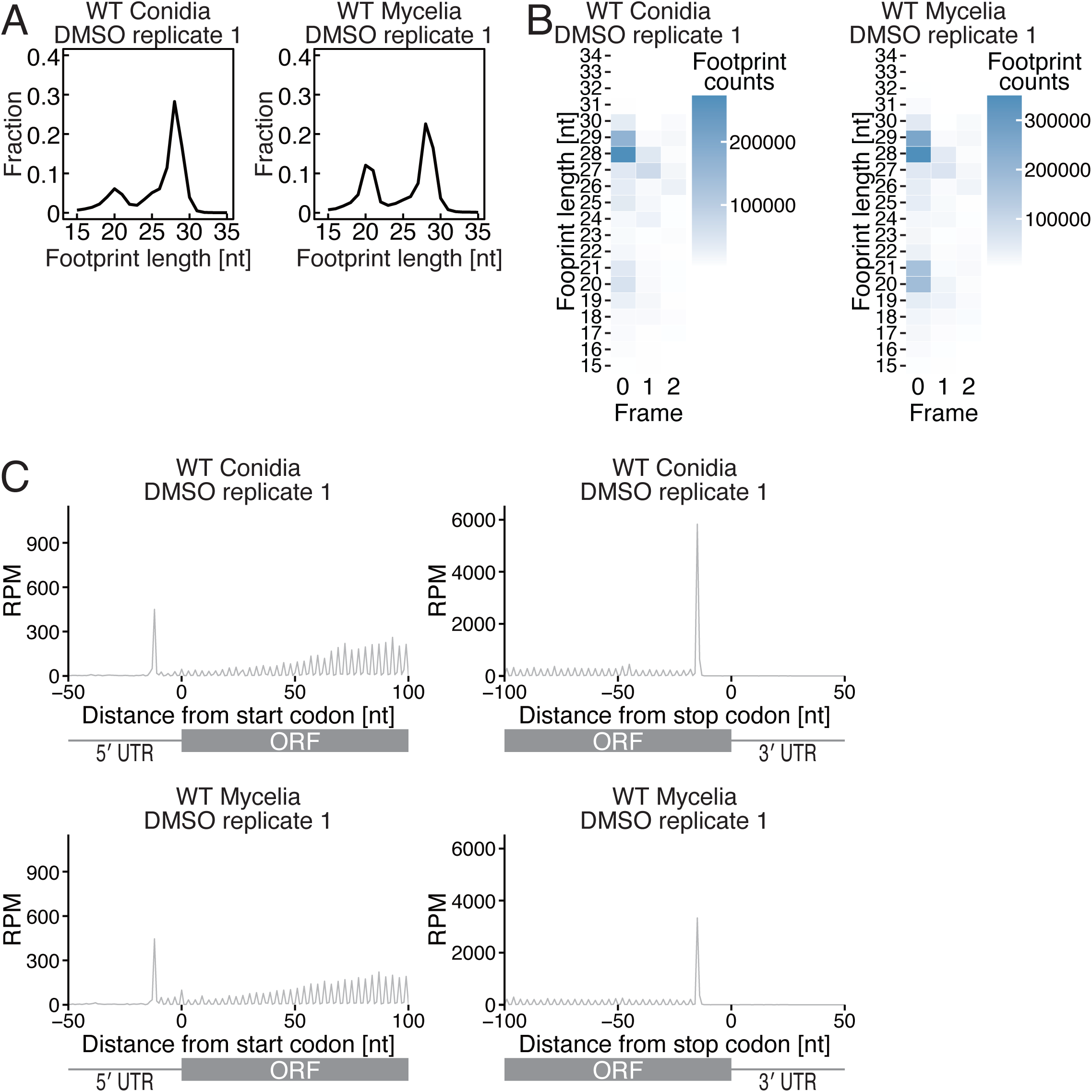
Characterization of ribosome footprints in *C. orbiculare*. (A) Distribution of ribosome footprint length in conidia and mycelia. (B) Tile plot of reading frames at each ribosome footprint length in conidia and mycelia. The 5′ end positions of the ribosome footprints are depicted. (C) Metagene plot of 29 nt ribosome footprints around start (left) and stop (right) codons in conidia and mycelia. The 5′ end positions of the ribosome footprints are depicted. RPM: Reads per million mapped reads.

**Figure 3 — figure supplement 3.**
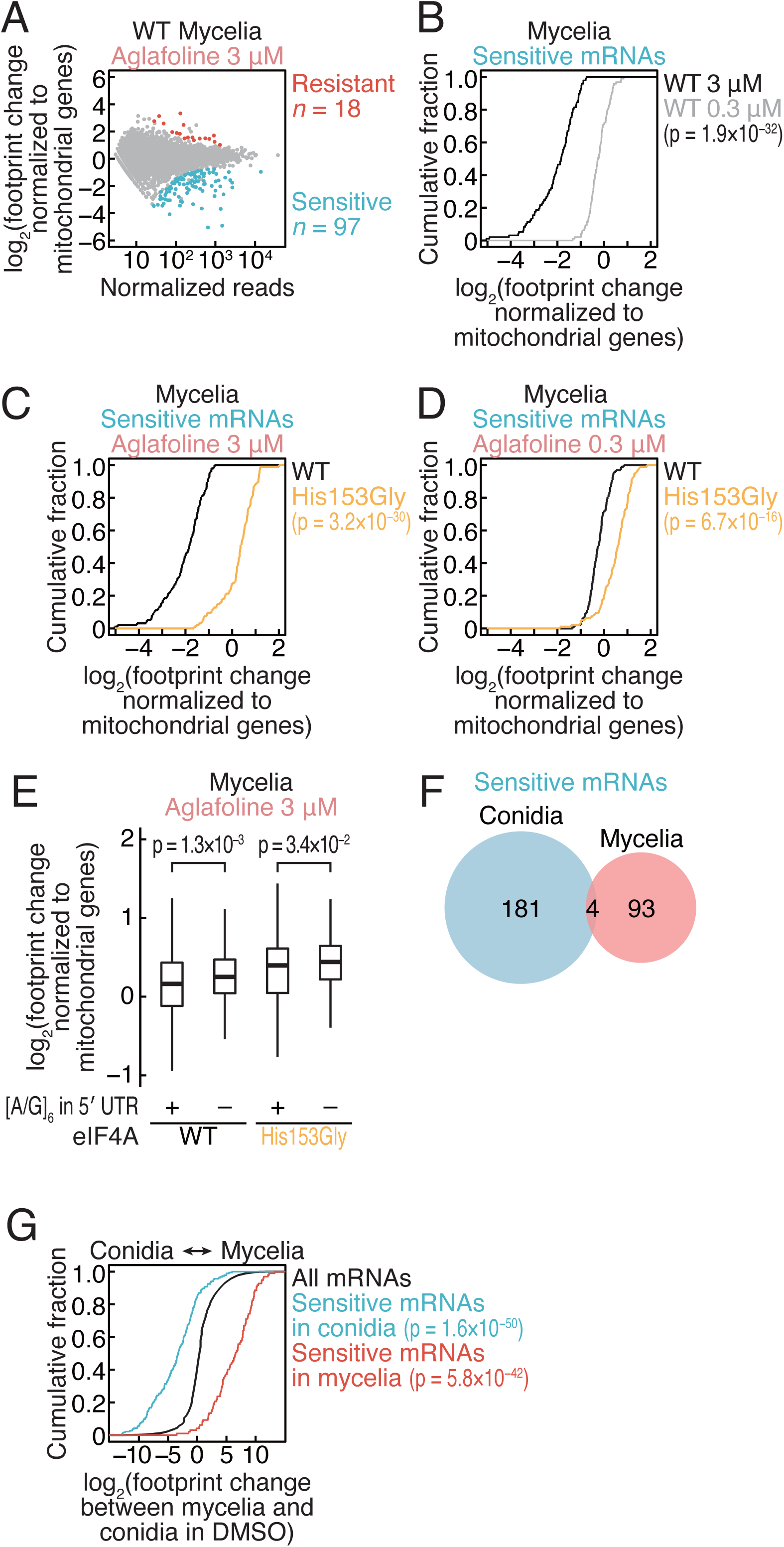
Translation changes by aglafoline treatment in recombined *C. orbiculare*. (A) MA plot of ribosome footprint changes by 3 µM aglafoline treatment in *C. orbiculare* eIF4A^WT^ mycelia. Resistant and sensitive mRNAs (false discovery rate [FDR] < 0.05) are highlighted. Data were normalized to mitochondrial footprints, which were used as internal spike-ins (Iwasaki et al., 2016). (B) Cumulative distribution of the ribosome footprint changes for aglafoline-sensitive mRNAs (defined in A) in *C. orbiculare* eIF4A^WT^ mycelia with 0.3 or 3 µM aglafoline treatment. (C and D) Cumulative distribution of the ribosome footprint changes for aglafoline-sensitive mRNAs (defined in A) by 3 µM (E) and 0.3 µM (F) aglafoline treatment in mycelia *C. orbiculare* eIF4A^WT^ and eIF4A^His153Gly^. (E) Box plot of ribosome footprint changes caused by 3 µM aglafoline treatment in mycelia across mRNAs with or without [A/G]_6_ motif in 5′ UTR. (F) Venn diagram of the overlap between aglafoline-sensitive mRNAs in conidia (defined in Figure 3B) and mycelia (defined in Figure 3 — figure supplement 3A). (G) Cumulative distribution of the ribosome footprint changes by the cell states (conidia biased as negative and mycelia biased as positive) for aglafoline-sensitive mRNAs in conidia (defined in Figure 3B) and mycelia (defined in Figure 3 — figure supplement 3A). The p values in B, C, D, E, and G were calculated by the Mann–Whitney *U* test.

**Figure 4 — figure supplement 1.**
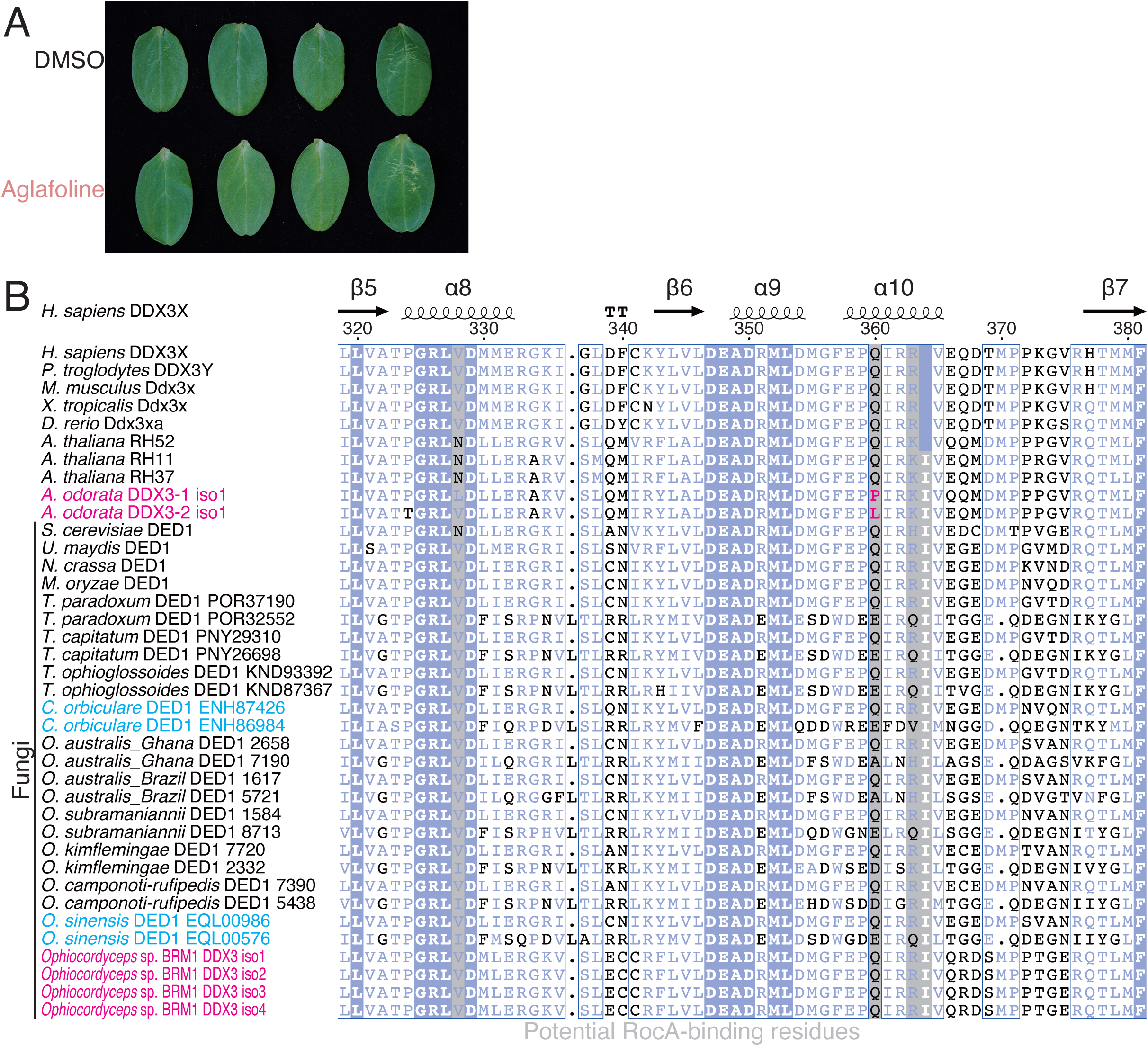
Characterization of cucumber leaves with aglafoline treatment. (A) *C. sativus* leaves were sprayed with DMSO or aglafoline (1 µM) in water and incubated for 3 d using the same method as *C. orbiculare* inoculation. (B) Alignment of DDX3 protein sequences from higher eukaryotes and fungal species, including *Ophiocordyceps* sp. BRM1 DDX3s.

**Supplementary Table 1. *De novo* assembly of the *Aglaia*-infecting fungus transcriptome.**

Summary of *de novo-*assembled transcripts and genes from *Aglaia*-infecting fungus RNA-Seq.

**Supplementary Table 2. Top 30 BLASTn hits of the *Aglaia*-infecting fungus ITS sequence against the NCBI nonredundant nucleotide database.**

Nucleotide sequence accessions were listed with subject strain, description, NCBI taxonomy ID, subject accession, and alignment statistics to *Aglaia*-infecting fungus ITS (percent identity, alignment length, mismatch numbers, gap opens, subject start, subject end, E-value, and bit score).

**Supplementary Table 3. List of fungal species for the multilocus phylogenetic tree analysis.**

Fungal species were listed with host species, strain names, GenBank IDs (ITS, SSU, LSU, *TEF1α*, and *RPB1*), and references. The DNA sequences shown in the columns are the best hits from the nucleotide collection searched by BLASTn.

**Supplementary Table 4. List of *C. orbiculare* strains used in this study.**

The *C. orbiculare* strains used in this study are listed with the strain IDs, genotypes, parental strains, and descriptions.

**Supplementary Table 5. List of oligonucleotides used in this study.**

The oligonucleotides used in this study are listed with the sequences, descriptions, and references.

**Figure 2 — figure supplement 1 — source data 1.**

Files for full and unedited gel images corresponding to Figure 2 — figure supplement 1B.

**Figure 3 — figure supplement 1 — source data 1.**

Files for full and unedited gel images corresponding to Figure 3 — figure supplement 1C.

